# A multimodal approach of microglial CSF1R inhibition and GENUS provides therapeutic effects in Alzheimer’s disease mice

**DOI:** 10.1101/2025.04.27.648471

**Authors:** Chinnakkaruppan Adaikkan, Md Rezaul Islam, P. Lorenzo Bozzelli, Matt Sears, Cameron Parro, Ping-Chieh Pao, Na Sun, TaeHyun Kim, Karim Abdelaal, Mia Sedgwick, Manolis Kellis, Li-Huei Tsai

**Affiliations:** Picower Institute for Learning and Memory, Massachusetts Institute of Technology, Cambridge, MA, USA; Department of Brain and Cognitive Sciences, Massachusetts Institute of Technology, Cambridge, MA, USA; MIT Computer Science and Artificial Intelligence Laboratory, Cambridge, MA, USA; Broad Institute of MIT and Harvard, Cambridge, MA, USA; Centre for Brain Research (CBR), Indian Institute of Science, Bangalore 560012, India

**Keywords:** Gamma oscillations, Microglia, PLX3397, GENUS, Alzheimer’s disease, MEF2C

## Abstract

The CSF1R inhibitor PLX3397, an FDA-approved treatment for a rare cancer, has been shown to reduce microglia count, lower inflammation, and increase synaptic markers in mouse models of Alzheimer’s disease (AD). However, the effects of PLX3397 on neural function in AD remain largely unknown. Here, we characterized the effects of PLX3397 treatment in 5xFAD mice. While PLX3397 increased synaptic density, it reduced the percentage of neurons phase-locked to gamma oscillations. This neural decoupling was closely associated with gene expression changes related to synapse organization. We investigated whether Gamma ENtrainment Using Sensory (GENUS) stimulation could counterbalance the neural circuit alterations induced by PLX3397. GENUS + PLX3397 restored gamma phase-locking, reshaped gene expression signatures, and improved learning and memory better than either treatment alone in 5xFAD mice. These findings suggest that CSF1R inhibitors like PLX3397 may benefit from a multimodal approach combining microglial targeting with non-invasive sensory stimulation to support neural physiology and improve cognitive function in AD.

## INTRODUCTION

Microglia are the resident macrophages of the brain involved in sensing and regulating neuronal activity^1^. Although microglial function is necessary for normal homeostatic brain function, aberrant activation is thought to drive neuroinflammation and the degeneration of synapses and neurons in Alzheimer’s disease (AD)^2^. During AD progression, microglia increase in numbers and demonstrate inflammatory phenotypes within most parts of the brain, including the cortex and hippocampus^3–7^. Furthermore, microglia impact the propagation of amyloid and tau pathology at various stages of disease progression^8–14^. Therefore, understanding the impact of altered microglial density and function in driving disease pathogenesis is of great therapeutic interest. Accordingly, pharmacological reduction of microglia via inhibition of colony-stimulating factor 1 receptor (CSF1R) – which is required for microglial survival – attenuates neuroinflammation and neurodegeneration in mouse models of AD^8–14^. However, despite numerous studies investigating the effects of depleting CSF1R-sensitive microglia on AD-associated pathological measures such as amyloid plaques, neuroinflammation, synaptic markers, and neurodegenerative phenotypes and gene expressions, there is limited understanding of how microglial depletion affects neural physiology and function *in vivo* in the context of AD. Recent studies that investigated the effect of systemic inhibition of CSF1R on neural oscillations have yielded conflicting results, with some studies indicating a lower threshold for seizure after CSF1R inhibitor treatments^15^, while others suggest an anti-epileptic effect of CSF1R inhibitor in rodent models^16^. Additionally, the CSF1R inhibitor PLX3397 (also known as pexidartinib) is approved by the Food and Drug Administration (FDA) for the treatment of adult patients with symptomatic tenosynovial giant cell tumors, a rare disease characterized by joint/soft tissue neoplasms^17,18^. Thus, understanding the impact of PLX3397 treatment on neural activity and function will provide insights into the impact of CSF1R-sensitive microglia on neural activity and will also instruct future treatment strategies for neurodegeneration and/or tumors.

Here, we observed that microglial depletion by chronic PLX3397 administration increased synaptic density and extracellular matrix without altering amyloid levels in the cortex of 12-month-old 5xFAD mice. However, PLX3397 administration negatively affected neural physiology, as evidenced by reductions in the number of neurons phase-locked to theta and gamma oscillations in the cortex. By treating the mice with non-invasive 40 Hz sensory stimulation (termed Gamma ENtrainment Using Sensory stimulation; GENUS), we were able to improve neural circuit function, promote phase-locking of neurons to gamma oscillations and improve novel object recognition memory in microglia-depleted 5xFAD mice. Molecular analysis revealed that entraining gamma frequency neural spiking by GENUS in these mice improved intrinsic neural mechanisms by notably enhancing the expression of MEF2C, a transcription factor critical to synaptic architecture and cognitive function^19,20^, as well as other genes involved in synaptic and extracellular matrix organization. This led to improved synaptic and extracellular architecture. These findings suggest that while targeting CSF1R can result in aberrant neural activity, this effect can be mitigated with a multimodal approach that promotes gamma oscillations in CSF1R-inhibited AD mice.

## RESULTS

### Chronic CSF1R inhibition with PLX3397 increases synaptic markers but disrupts neuronal synchronization in 5xFAD mice

To study how CSF1R-sensitive microglia impact neuropathology and neural activity in AD, we subjected 11-month-old 5xFAD mice, one of the most commonly used amyloidosis mouse models of AD^21^, to a diet containing the CSF1R inhibitor PLX3397 for 50 days and performed immunohistochemical (IHC) analysis and electrophysiological recordings. We used the CSF1R inhibitor PLX3397 at a concentration of 600 ppm, which was previously shown to reduce microglia *in vivo* effectively^22^. The control group was age-matched 5xFAD littermates that received a regular control diet. We first verified the effect of chronic PLX3397 on amyloid and synaptic levels in 5xFAD mice (**Figure 1A-1D**). IHC analysis revealed that chronic PLX3397 administration did not affect amyloid levels (arbitrary units (au), 9894 ± 1181) as analyzed by the D54D2 positive amyloid signal in the visual cortex compared to control diet mice (7795 ± 762.5) (**Figure 1A**). This observation is consistent with previous findings in 5xFAD mice^10,23^. Next, we sought to replicate key findings of earlier studies that showed that chronic PLX3397 administration led to higher levels of synaptic markers. Indeed, PLX3397 administration impacted synaptic integrity, manifesting as a reduction in C1q (**Figure 1B**) and a concomitant increase in synaptophysin signals, which are markers of synaptic pruning and integrity, respectively (**Figure 1C**). The higher levels of synaptic markers after chronic PLX3397 administration are consistent with recent reports^10,13^, suggesting the potential of PLX3397 to increase synaptic markers in a model of AD. These findings prompted us to investigate the direct effects of PLX3397 administration on neuronal integrity. Perineuronal nets (PNNs), specialized extracellular matrix structures, are known to regulate the activity of neurons, with microglia playing a crucial role in their organization^11,24–26^. Notably, we observed that PLX3397 administration led to an increase in WFA+ (*Wisteria floribunda agglutinin*) labeling of PNNs, indicating enhanced PNN signals (**Figure 1D**). Collectively, these findings highlight the significant impact of CSF1R inhibition on neuronal architecture and synaptic density.

**Figure 1.**
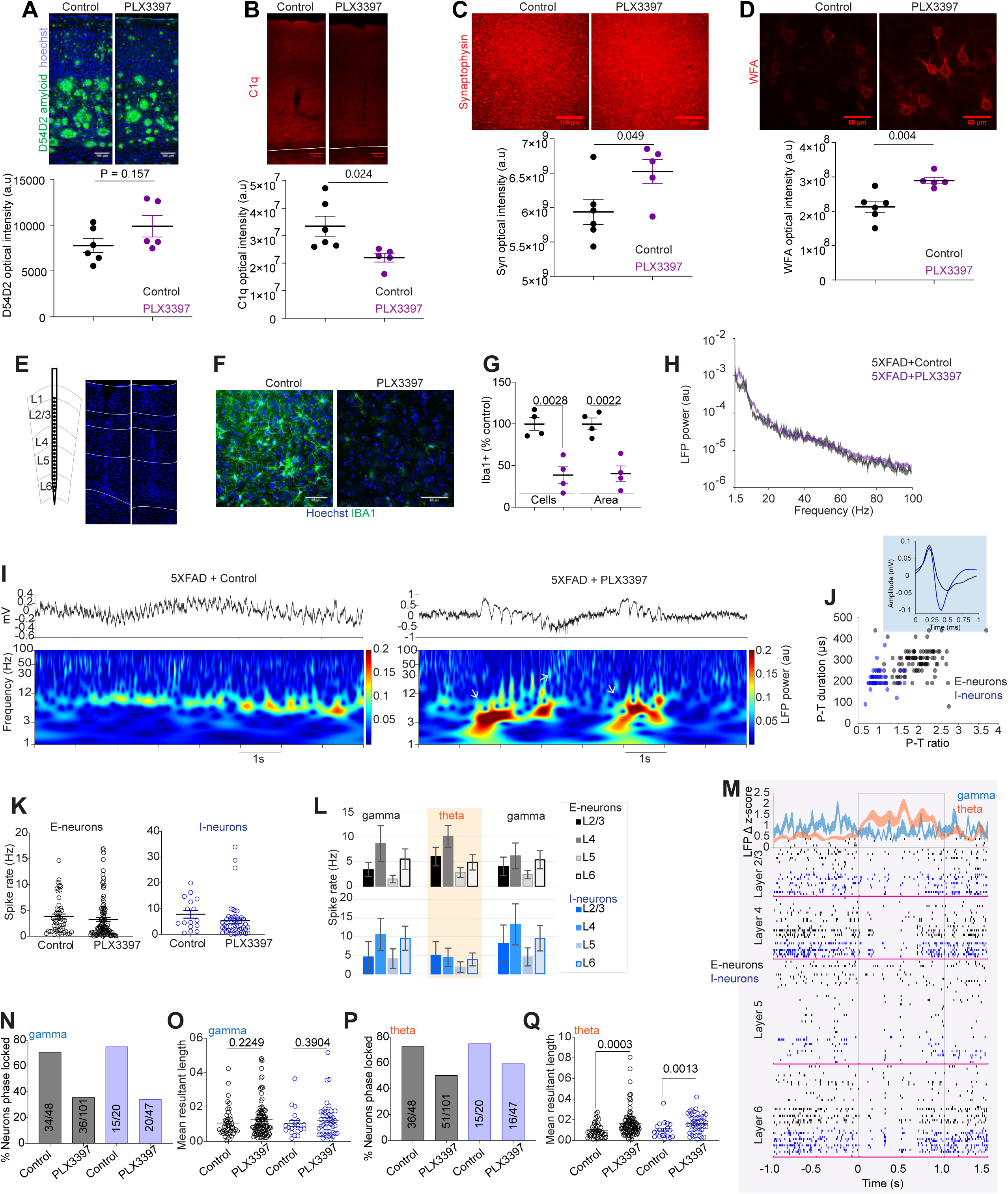
Chronic PLX3397 treatment reduces the percentage of gamma phase-locked neurons in 5xFAD mice. (A) Confocal images show D54D2-positive amyloid and Hoechst. PLX3397 did not affect amyloid (unpaired, two-tailed t-test, t = 1.544, P = 0.157). n = 5-6 mice per group. (B) Confocal images show the complementary molecule C1q. PLX3397 reduced C1q levels (t = 2.699, P = 0.0244). (C) Confocal images show synaptophysin. Plx3397 increased synaptophysin (t = 2.273, P = 0.049). (D) Confocal images show *Wisteria floribunda* agglutinin (WFA). PLX3397 increased WFA signals (t = 3.774, P = 0.004). (E) *In vivo* electrophysiological recording configuration. Linear probes were implanted in the visual cortex. Example images of control (left) and PLX3397 (right) mice show linear probe recording locations (Hoechst 3352 stain, blue). (F) Confocal images show IBA1 in control and PLX3397-treated 5xFAD mice. (G) The number of IBA1+ cells (unpaired, two-tailed t-test, t = 4.879, P = 0.0028) and IBA1+ area (unpaired, two-tailed t-test, t = 5.092, P = 0.0022) were reduced after PLX3397 administration in 5xFAD mice (n = 4 mice/group). (H) LFP Power in control and PLX3397-treated 5xFAD mice. 2W RM ANOVA, treatment x frequency interaction, F (201, 1206) = 2.519, P < 0.0001. There was no group difference between control and PLX3397-treated mice (2W RM ANOVA, F (1, 6) = 3.425, P = 0.0993). (I) Plots show unprocessed raw LFP traces and the corresponding time-resolved LFP power from 5xFAD without or with PLX3397 administration. (J) The plot shows the duration and ratio of peak-to-trough (P-T) of single units from PLX3397 5XFAD mice. Inset, spike waveforms of representative E-neuron and I-neuron (extracellular spike waveforms were inverted for illustrative purposes). (K) The mean spike rate of E-neurons (unpaired, two-tailed t-test, t = 0.9159, P = 0.3612) and I-neurons (unpaired, two-tailed t-test, t = 1.273, P = 0.2077) did not differ between control and PLX3397-treated 5xFAD mice. (L) The mean spike rate of neurons in each cortical layer at the onset (0 to −200 ms) of higher theta-burst activity and ±800 ms. (M) Single-unit raster plot shows spiking during pre-, post-, and theta-bursts in PLX3397-treated 5xFAD mice. (N and O) Plots show the percentage of E-neurons (gray) and I-neurons (blue) phase-locked to gamma oscillations (N) and the strength of phase locking (O) in PLX3397 and control 5xFAD mice. (P and Q) Plots show the % of phase-locked neurons to theta (P) and the strength of phase locking (Q). Numbers in charts N and P represent neurons out of the total neurons significantly phase-locked (Rayleigh statistics, P < 0.05) to gamma and theta oscillations in each comparison. Scale bar = 100 (A, B, C) or 50 μm (D, F) as indicated. au = arbitrary units.

We next asked whether the increased levels of synaptic markers and extracellular matrix proteins impacted neural oscillatory activities. To address this, we performed high-density linear probe recordings in awake 5xFAD mice after 50-day administration of PLX3397. Post-recording, we verified the recording site in the visual cortex by histology (**Figure 1E**) and evaluated PLX3397-associated microglia reduction by assessing IBA1+ cells. The PLX3397 treatment significantly reduced microglia number (100 ± 7.64 versus 38.57 ± 10.0 IBA1+ cells in the control and PLX3397 diet, respectively) and area (100 ± 7.14 versus 40.34 ± 9.28) per region of interest (ROI) (**Figure 1F and 1G**).

To explore the effect of PLX3397 administration on neural oscillations, we examined the local field potential (LFP) power. We observed an overall LFP power change as a function of frequency in both groups; however, there was no significant difference in the PLX3397 group compared to the control 5xFAD group (**Figure 1H**). Interestingly, time-resolved analysis of the LFP revealed that chronic PLX3397 administration results in elevated power of gamma and theta that occur incongruently (**Figure 1I; Figure S1A**). Specifically, PLX3397-treated 5xFAD mice exhibit an alternation of gamma (30-50 Hz) and theta (3-12 Hz) (**Figure S1A and S1B**). Conversely, control 5xFAD mice did not demonstrate this alternating oscillatory state (**Figure 1I; Figure S1A - S1C**). The multiunit activity across all cortical layers showed marked differences between control and PLX3397-administered 5XFAD mice (**Figure S1D**). These observations led us to study the relationships between the aberrant LFP oscillations and neuronal spiking patterns after PLX3397 administration. To this end, we isolated single units and classified them into either putative excitatory neurons (E-neurons) or interneurons (I-neurons) (**Figure 1J**) using the parameters as described previously^27^. We isolated 47 and 102 E-neurons and 20 and 47 I-neurons from the control and PLX3397 groups, respectively. We did not detect differences in the overall mean spiking rate of E-neurons (control, 3.84 ± 0.49; PLX, 3.23 ± 0.39) and I-neurons (control 7.87 ± 1.51; PLX, 5.38 ± 1.01) (**Figure 1K**). However, the spiking patterns in the PLX3397 group markedly differed between LFP high theta activity and gamma (**Figure 1L**). As shown in **Figure 1M** single unit raster plots and the aggregated plots (**Figure 1L; Figure S1E**), spiking rates of I-neurons during high theta activity were significantly lower than those of the gamma. This reduction was evident across all cortical layers during theta (**Figure 1L, 1M**). As theta activity subsides and gamma emerges, neuronal spiking increases significantly (**Figure 1L; Figure S1ED**). At the population level, E-neurons maintained their spiking rates until the onset of the theta; I-neurons reduced their overall spiking rate preceding and during the theta activity (**Figure 1L and 1M; Figure S1E**). We observed that layer 4 (L4) E-neurons transiently increased their spiking closer to the onset of the theta activity (**Figure 1L; Figure S1E**). In contrast, I-neurons in layers 4 and 6 exhibited higher spiking during gamma (**Figure 1L**). These data suggest that neurons alter their spiking patterns between LFP theta activity and gamma rather than simply changing their overall mean spiking rate following PLX3397-mediated depletion of CSF1R-sensitive microglia.

Given that neuronal phase-locking to oscillatory activity is crucial for memory and cognitive function^28^, we next assessed the phase-locking of neurons to explore the relationship between neuronal spiking and LFP oscillations after PLX3397 treatment. We quantified the percentage of LFP phase-locked neurons and the strength of phase locking using Rayleigh statistics and mean resultant vector length, respectively. These analyses revealed that the percentage of both E-neurons and I-neurons in the PLX3397 group were less phase-locked to 30-50 Hz gamma oscillations than neurons in the control 5xFAD group (**Figure 1N**), without significant difference in the strength of those phase-locked neurons to gamma oscillations (MRL, E-neurons; control, 0.106 ± 0.012; PLX, 0.126 ± 0.010. I-neurons; control, 0.105 ± 0.017; PLX, 0.123 ± 0.014) (**Figure 1O**). Furthermore, we observed that PLX3397 treatment significantly affected both the percentage of E-neurons and I-neurons phase-locked to 3-12 Hz theta oscillations (**Figure 1P**) as well as the strength of theta phase-locking of neurons (**Figure 1Q**), manifesting as a reduction in the percentage of phase-locked neurons and an increase in the phase-locking strength (MRL, E-neurons; control, 0.094 ± 0.009; PLX, 0.163 ± 0.012. I-neurons; control, 0.101 ± 0.017; PLX, 0.165 ± 0.013).

These observations suggest that while CSF1R-sensitive microglia inhibition increased synaptic densities and neuronal architecture, it also disrupts gamma oscillations *in vivo*, reducing the number of neurons phase-locked to gamma oscillations. Given these effects, we next investigated whether entraining neural spiking and oscillations at gamma frequency could reverse the PLX3397-induced alterations in neural oscillations in 5xFAD mice. This question is particularly important, as PLX3397 has been FDA-approved and may have significant implications for AD-associated pathologies.

### GENUS improves neural activity in PLX3397-treated 5xFAD mice

We reasoned that evoking gamma oscillations, which are thought to be modulated by PV+ interneurons^29^, could induce I-neuronal activity to improve the neuronal phase-locking and aberrant neural activity caused by PLX3397 administration. To answer this, we exposed PLX3397-treated 5xFAD mice to 40 Hz visual gamma stimulation and found that it robustly induced 40 Hz LFP spectral power (**Figure 2A and 2B**). Thus, CSF1R-sensitive microglial depletion did not abolish the ability of mice to entrain patterned gamma frequency (i.e., 40 Hz) visual stimuli. We observed that acute 40 Hz visual stimulation induced physiological LFP waveforms and reduced the aberrant theta activity (PLX, 3.043 ± 0.132 s; PLX+40 Hz, 4.547 ± 0.513 s) (**Figure 2B; Figure S2A**). Interestingly, PLX3397-administered 5xFAD mice exhibited greater power of 40 Hz entrainment compared to control 5xFAD mice (**Figure S2B and S2C**). Together, these observations suggest that gamma stimulation can counteract the dysregulated neural oscillations induced by CSF1R-sensitive microglial depletion and that their combined effects produce a more robust entrainment response.

**Figure 2.**
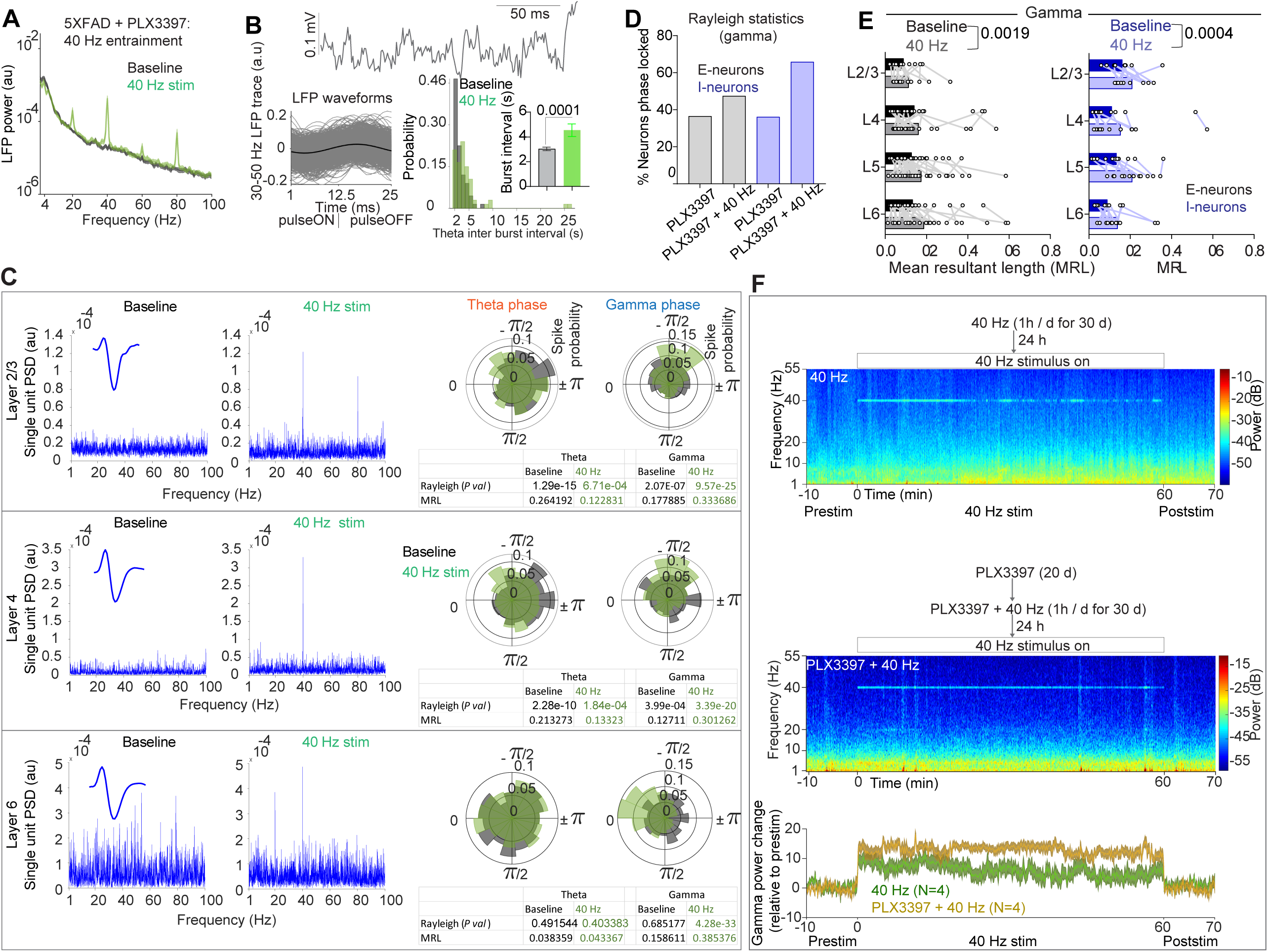
GENUS improves neural function in PLX3397-treated 5xFAD mice. (A) LFP power during baseline and 40 Hz stimulation. n = 4 mice/group, au = arbitrary units. (B) Top: LFP trace during 40 Hz entrainment. Bottom left: LFP waveforms as a function of 40 Hz stimulus. Bottom right: aberrant theta activity was significantly reduced during acute 40 Hz entrainment in PLX3397-treated 5xFAD mice (Mann-Whitney U = 1304, P = 0.0001). (C) Three simultaneously recorded I-neurons from L2/3, L4 and L6 that showed 40 Hz entrainment. Polar plots (right) show spike probability along LFP theta and gamma phases during baseline and 40 Hz entrainment. Rayleigh statistics and mean resultant length (MRL) indices indicate whether neuronal spiking is phase-locked to LFP and the phase-locking strength, respectively. (D) A higher percentage of E-neurons and I-neurons were phase-locked to gamma during 40 Hz gamma entrainment. (E) Phase-locking strength of E-neurons (2W ANOVA, F (1, 97) = 10.21, P = 0.0019) and I-neurons (2W RM ANOVA, F (1, 43) = 14.50, P = 0.0004) to LFP gamma were higher during 40 Hz entrainment. Data are single units. (F) Experiment outline (top). LFP power spectrogram before, during, and after 40 Hz stimulation in GENUS and PLX3397+GENUS (middle) treated 5xFAD mice. The line plot (bottom) shows the gamma power relative to baseline.

To determine whether the reduction in aberrant oscillatory activity in PLX3397-treated 5xFAD mice was due to rhythmic spiking of neurons during 40 Hz entrainment, we analyzed neuronal spiking rhythmicity and phase locking to LFP oscillations during acute 40 Hz entrainment. Among the 47 recorded I-neurons, 13 neurons exhibited 40 Hz entrainment, as indicated by a distinct 40 Hz peak in the power spectral density of units’ activity (**Figure 2C**). However, across these 13 40 Hz rhythmic I-neurons, we could not establish whether the spiking occurred during any specific phase of LFP gamma (**Figure 2C**). Interestingly, despite only a subset of I-neurons (13 of 47) entraining at 40 Hz, phase-locking analysis revealed that 40 Hz entrainment dramatically increased the percentage of I-neurons phase-locked to gamma (65.95% versus 34.04% at baseline) (**Figure 2D**). Although we did not identify any 40 Hz rhythmic E-neurons, we observed a marked increase in the percentage of LFP gamma phase-locked E-neurons during 40 Hz stimuli (46.53% versus 35.64% in the baseline) (**Figure 2E**). We next examined whether neurons in specific layers of the cortex entrain 40 Hz and whether they are phase-locked. Overall, we observed that I-neurons in L2/3, L4, and L6 show 40 Hz entrainment (**Figure 2C**). Furthermore, both E-neurons and I-neurons distributed across all layers of the cortex (E-neurons, baseline Vs 40 Hz, L2/3, 0.090 ± 0.014 Vs 0.115 ± 0.021; L4, 0.142 ± 0.033 Vs 0.162 ± 0.031; L5, 0.128 ± 0.013 Vs 0.175 ± 0.020; and L6, 0.134 ± 0.021 Vs 0.189 ± 0.0219. I-neurons, L2/3, 0.162 ± 0.024 Vs 0.210 ± 0.021; L4, 0.111 ± 0.043 Vs 0.154 ± 0.048; L5, 0.134 ± 0.028 Vs 0.209 ± 0.023; and L6, 0.090 ± 0.013 Vs 0.139 ± 0.025) showed enhanced gamma phase-locking strength (**Figure 2E**). The theta phase-locked E-neurons but not I-neurons were modestly reduced during 40 Hz entrainment without a significant difference in the strength of the theta phase-locking of neurons (**Figure S2D and S2E**). Together, these results suggest that acute 40 Hz sensory stimulation (i.e., GENUS)^30,31^ induces 40 Hz rhythmic spiking of I-neurons in many cortical layers and enhances the percentage of neurons phase locked to gamma oscillations and further reduces the aberrant alterations of oscillations caused by PLX3397 treatment.

Although 40 Hz entrainment was evident during acute sensory stimulation, we sought to determine whether chronic GENUS could sustain entrainment in both control and PLX3397-treated 5xFAD mice. To this end, we subjected control and PLX3397-treated 5xFAD mice to 40 Hz stimulation one hour per day for 30 days and then performed electrophysiological recordings. We observed that 40 Hz stimulation robustly induced 40 Hz entrainment across the entire one-hour stimulation period in these mice (**Figure 2F**), and I-neurons in L2/3, L4 & L6 in the chronic PLX3397+GENUS group were 40 Hz rhythmic (**Figure S2F**). Interestingly, mice treated with the combination of the PLX3397 diet and GENUS exhibited a stronger 40 Hz entrainment than mice that received GENUS alone (**Figure 2F**).

In the next set of biochemical and molecular analyses, we characterized how PLX3397 administration alone impacted the cellular and molecular functions associated with aberrant neural activity and how GENUS countered these to improve neural network functions.

### Reduced inflammation and improved myelination after combined PLX3397 and GENUS

First, previous studies have shown that PLX3397 and chronic GENUS independently reduce microglial density and modulate neuroinflammation in multiple mice models^10,32^. Thus, we investigated the combined effects of PLX3397 and GENUS on microglial density, inflammatory markers, and gene expressions. We treated 5xFAD mice with PLX3397 via oral delivery in mouse chow. After 20 days of administration of PLX3397, both untreated- and PLX3397-treated 5xFAD mice were subjected to 30 days of daily GENUS (PLX3397-treated mice were maintained on PLX3397 for a total of 50 days) (**Figure 3A**). Following completion of the treatments, we quantified microglial density per ROI in four groups: 1) untreated (control), 2) PLX3397 alone, 3) GENUS alone, and 4) combined PLX3397+GENUS-treated 5xFAD mice (**Figure 3A and 3B**). We observed a significant reduction in IBA1+ microglia number in PLX3397 (microglia number, 47.18 ± 5.44), and PLX3397+GENUS (24.34 ± 2.41) groups compared to controls (100 ± 5.12) (**Figure 3B and 3C**). Notably, the GENUS-only group also showed a reduction in the IBA1+ cells (84.0 ± 6.42 & 9.17 ± 0.85) compared to controls (**Figure 3C**), which is consistent with our previous findings in CK-p25 mice^30^.

**Figure 3.**
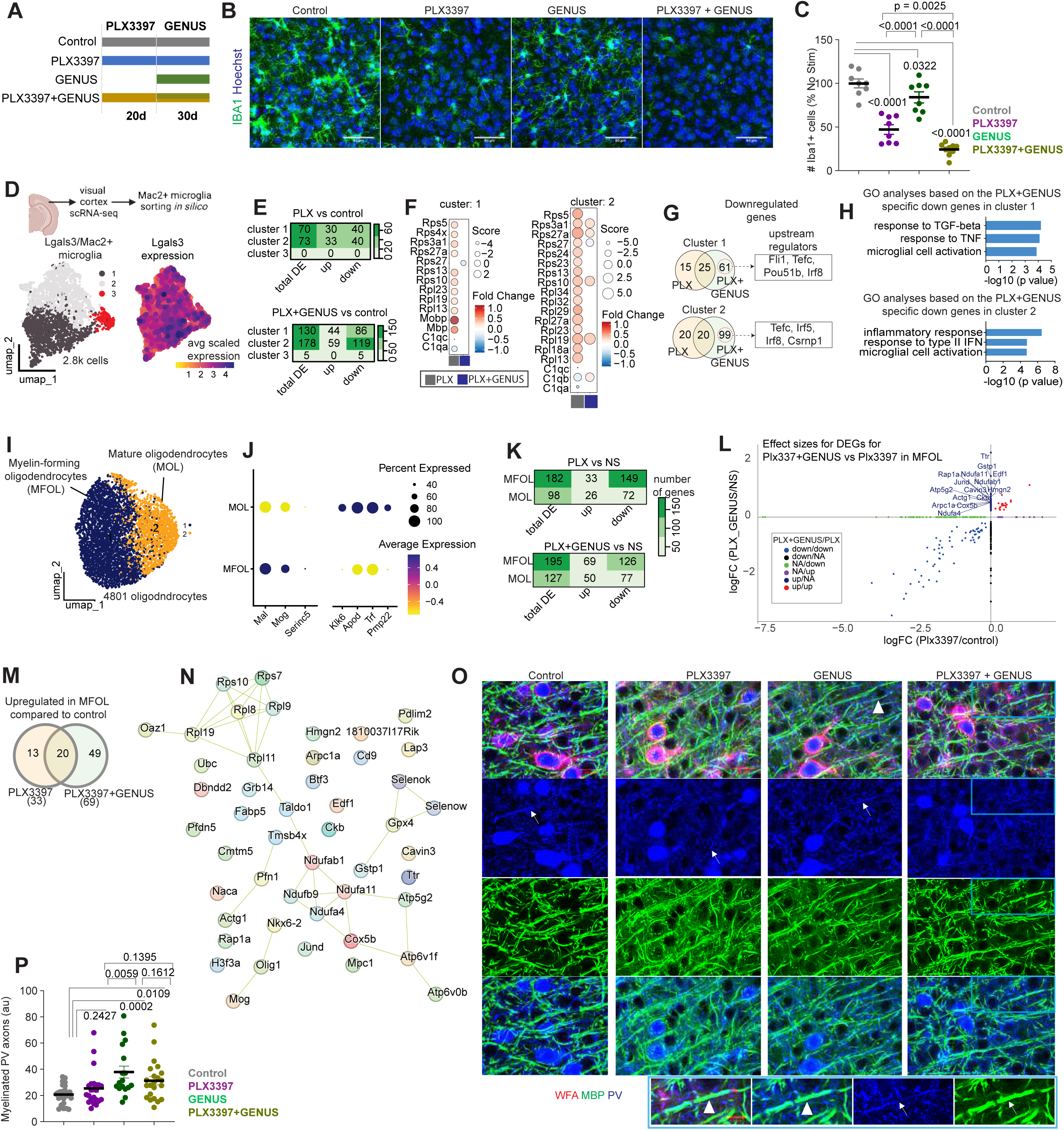
Chronic GENUS improves myelination in PLX3397-treated 5xFAD mice. (A) Experiment outline to administer CSF1R inhibitor and/or GENUS in 5xFAD mice. (B) Confocal images show IBA1. Scale bar = 50 μm. (C) IBA1+ microglial numbers (n = 8 - 9 mice/group; AONVA, F (3, 29) = 49.37, P = 0.0001) expressed as % no treatment control. (D) UMAP visualization of Lgals3/Mac2+ microglia clusters based on single-cell RNA-seq data from the visual cortices of 5xFAD mice. Mac2+ cells were sorted *in silico* before unbiased clustering. (E) Differentially expressed genes (DEGs) across clusters in the PLX (PLX3397) and PLX+GENUS (PLX3397+GENUS) groups, compared to the control group. The color scale indicates the number of DE genes. (F) DEGs name in clusters 1 and 2. (G) Venn diagram shows the overlap of downregulated genes in clusters 1 and 2 between the PLX vs. control and PLX+GENUS vs. control comparisons. Boxes highlight the top upstream regulators for differentially downregulated genes specific to the PLX+GENUS treatment. (H) Gene Ontology (GO) analysis of the PLX+GENUS-specific downregulated genes in clusters 1 and 2. (I) UMAP clustering of 4.8k oligodendrocytes based on single-cell RNA-seq data revealed two distinct clusters: Myelin-forming oligodendrocytes (MFOL) and mature oligodendrocytes (MOL). (J) Enrichment of cell-type-specific markers in the two types of oligodendrocytes. Dots represent the percentage of cells expressing each marker, and the color scale indicates the scaled average expression level. (K) Heatmaps show DEGs in MFOL and MOL following PLX3397 and PLX3397+GENUS treatments. Differential expression was determined by comparing each treatment group to the control condition. (L) Comparison of effect sizes for differentially expressed genes between the PLX3397+GENUS vs PLX3397 groups in MFOL. The y-axis represents the log fold change (logFC) of genes in the PLX3397+GENUS vs. control comparison, while the x-axis represents the logFC of genes in the PLX3397 vs. control comparison. (M) Venn diagram shows the overlap of upregulated genes in MFOL between PLX3397 and PLX3397+GENUS conditions, both compared to the control treatment. Forty-nine genes were uniquely upregulated following chronic GENUS treatment compared to the control condition. (N) Network plot highlighs genes uniquely upregulated in MFOL after chronic GENUS treatment in PLX3397-treated 5xFAD mice. Increased expression of transcription factors related to oligodendrocyte differentiation and myelination, such as Mog, Olig1, and Nkx2-2, were observed in the myelin-forming oligodendrocytes. (O) Co-labeled WFA, MBP, and PV confocal images are shown (scale bar = 20 μm; inset 10 μm). (P) The summary graph shows myelinated PV axons (ANOVA F (3, 83) = 5.895, P = 0.0011).

Next, we assessed the functional status of microglia by examining their transcriptomic signatures. To this end, we profiled 27,607 high-quality single cells by single-cell RNA sequencing (scRNA-seq) from the visual cortices, representing various glial and vascular cell types, including microglia, oligodendrocytes, meningeal fibroblasts, endothelial cells, T and mural cells (**Figure S3**). We first focused on identifying transcriptional changes in Mac2+/Lgals3+ PLX3397-resistant microglia^33^. We sorted Mac2+/Lgals3+ microglia *in silico* and identified three distinct transcriptional subtypes (**Figure 3D**). Differential gene expression analyses revealed that PLX3397 treatment led to a comparable number of gene expression changes in clusters 1 and 2 but not in cluster 3 (**Figure 3E**). Next, we focused our downstream analyses on clusters 1 and 2, where we observed a relatively higher number of downregulated genes in both clusters after treatment (**Figure 3F, Table S1**). We then examined the upstream regulators of genes uniquely downregulated in the PLX3397+GENUS group (**Figure 3F**). In cluster 1, the top upstream regulators included *Fli1, Tefc, Pou51b, Irf8*, while in cluster 2, they included *Tefc, Irf5, Irf8, Csrnp1*. These regulators are commonly associated with inflammatory responses (**Figure 3G**). Consistent with this, gene ontology (GO) analysis of the downregulated genes specific to the PLX3397+GENUS group revealed several inflammatory processes, including microglial cell activation (in both clusters 1 and 2), response to tumor necrosis factor (cluster 1), and regulation of inflammatory signaling pathways (such as TGF-β and IFNs) (**Figure 3H**). These observations of reduced inflammation in microglia after Plx3397+GENUS treatment are interesting because a reduction in neuroinflammation can be considered beneficial in AD. Moreover, neuroinflammation induced by lipopolysaccharide injection has been shown to affect PV interneurons and neural activity in mice^34^. Similarly, *ex vivo* experiments showed that IFN induced microgliosis and inflammation affected gamma oscillations^35^, Therefore, it is reasonable that the improved network function in PLX3397-treated 5xFAD mice after GENUS is associated with the reduction in neuroinflammation. Concomitant with the reduced inflammation, we considered that PLX3397+GENUS administration could also impact cellular and molecular functions of neurons.

Neuronal functions, especially parvalbumin (PV) interneurons^36^, are regulated by oligodendrocytes through myelination. PV interneurons are shown to be participating in light flicker-evoked gamma ^37^. Therefore, we investigated the effects of the treatment conditions on oligodendrocytes in our scRNA-seq data. We categorized oligodendrocytes into myelin-forming and mature oligodendrocyte subsets expressing unique marker genes (**Figure 3I and 3J**). Differential expression analyses revealed that both PLX3397 and PLX3397+GENUS treatments induced more differentially expressed genes (DEGs) in myelin-forming oligodendrocytes compared to mature oligodendrocytes, relative to controls (**Figure 3K, Table S2**). Given the stronger transcriptional response in myelin-forming oligodendrocytes, we focused on this subset and identified genes uniquely upregulated in PLX3397+GENUS-treated mice, particularly those involved in lipid metabolism (*Gpx4, Fabp5*) (**Figure 3L-3N**). Maintenance of myelin requires continuous lipid turnover^38^ and constant high-level expression of myelin proteins^39^. Moreover, myelinating oligodendrocytes support fast-spiking axons by providing lactate or pyruvate^40^ for the generation of ATP^41^. Consistent with this, PLX3397+GENUS treatment upregulated genes linked to mitochondrial function and oxidative phosphorylation including subunits of the NADH dehydrogenase (*Ndufab1, Ndufa4, Ndufa11, Ndufb9*), subunit of cytochrome c oxidase (*Cox5b*), subunits for ATP synthase (*Atp5g2, Atp6v1f, Atp6v0b*), mitochondrial pyruvate carrier (*Mpc1*), and gene involved in ATP buffering (Creatine kinase B-type, *Ckb*). Additionally, genes involved in the pentose phosphate pathway (*Taldo1*) and redox signaling (*Selenow, Selenok*) were upregulated. Furthermore, genes involved in protein folding and degradation (e.g., *Ubc, Pfdn5*) and gene transcription regulation (e.g., *Jund, Hmgn2*) were also upregulated (**Figure 3N**) in the PLX3397+GENUS group. Notably, PLX3397+GENUS treatment significantly upregulated transcription factors (*Nkx6-2, Olig1*) essential for oligodendrocyte differentiation and myelination (**Figure 3N**). Additionally, Mog expression, which encodes myelin oligodendrocyte glycoprotein, a marker for oligodendrocyte maturation^42^, was significantly increased. These results align with recent studies demonstrating that 40 Hz sensory stimulation promotes myelination in a mouse model of demyelination^43^ and chemobrain^44^.

To further validate these findings, we performed additional IHC analysis. GENUS treatment increased myelin ensheathment of axons of PV interneurons in PLX3397-treated 5xFAD mice (20.75 ± 1.45, 25.44 ± 2.83, 37.91 ± 4.56, & 31.3 ± 3.21 in control, PLX3397, GENUS, PLX3397+GENUS groups, respectively) (**Figure 3O, 3P; Figure S7C**). Thus, repeated GENUS treatment enhances axonal myelination of PV interneurons in PLX3397-treated 5xFAD mice.

### Improved MEF2C target genes and synapse organization in 5xFAD mice following combined PLX3397 and GENUS treatment

We next asked how regulating gamma oscillations affects synaptic and neural circuit function in PLX3397-treated 5xFAD mice. To investigate this, we employed an unbiased RNA sequencing approach. Because scRNA-seq did not capture neuronal populations (**Figure S3**), we performed snRNA-seq to better profile neuronal transcriptomes. We generated and analyzed high-quality transcriptomes from 33,112 nuclei derived from the visual cortex (**Figure 4A; Figure S4**). As shown in **Figure S4,** these nuclei represented various cell types. The majority of the cells captured were excitatory and inhibitory neurons (**Figure 4A and Figure S4**). PLX3397+GENUS versus PLX3397 comparison revealed several DEGs across cell types. Notably, the excitatory neuron cluster (E-neuron) exhibited the highest number of DEGs (**Figure 4B; Table S3**). We focused on differentially expressed genes in neurons for downstream analyses. GO analyses revealed that the upregulated genes after PLX3397+GENUS treatment in E-neurons were associated with synaptic plasticity (*Nrgn, Cplx2, Cfl1*), cognition (*Mef2c, Ube3a*), and general synaptic organization and function (Sparcl1, Ube3a, Apoe) (**Figure 4B and 4C**). Similarly, upregulated genes related to synapse organization and synaptic plasticity were also evident in the inhibitory neurons (I-neurons) (**Figure 4B and 4C; Figure S5**). Upregulated genes enriched for RNA splicing and cognition were exclusively observed in E-neurons (**Figure 4B and 4C**).

**Figure 4.**
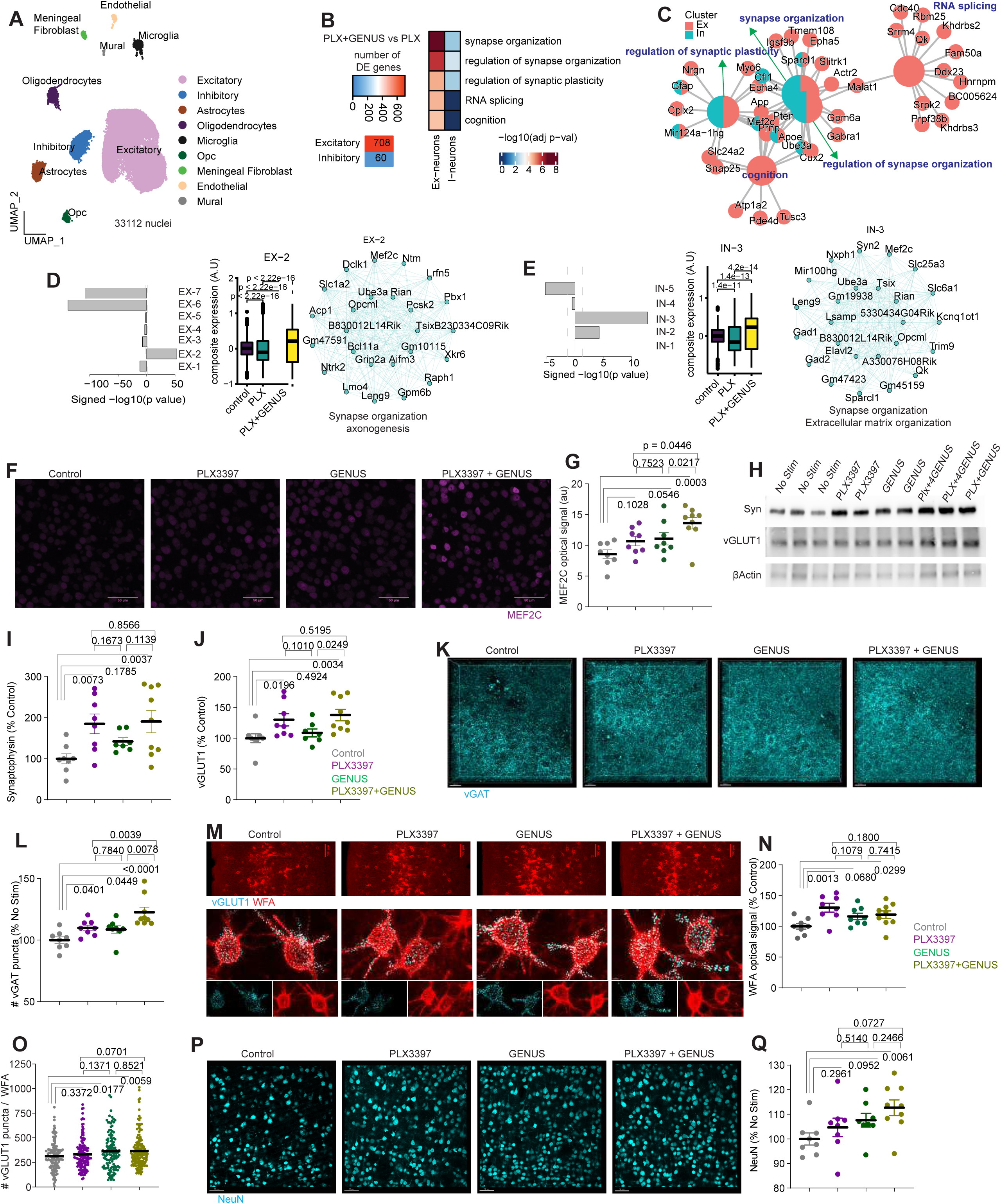
Chronic GENUS enhances MEF2C in PLX3397-treated 5xFAD mice. (A) UMAP visualization of snRNA-seq colored by cell type. (B) Left: heatmap shows differentially expressed genes in the PLX+GENUS group compared to PLX alone across various cell types. Right: significant biological processes of the upregulated genes from excitatory and inhibitory neurons. (C) Network map shows genes enriched for the biological processes. Smaller nodes represent genes, while the larger nodes represent the gene ontology terms. Nodes with shared color highlight enrichment in both excitatory and inhibitory neurons. (D) Excitatory gene modules based on gene co-expression patterns. Top: bar plot shows the significance of the excitatory gene modules and standardized mean difference between PLX+GENUS and PLX alone groups. Bottom: boxplot shows the composite expression of module Ex-2 among control, PLX+GENUS and PLX groups. Right panel: gene network displays the top 25 hub genes from the Ex-2 module. (E) Left: bar plot displays modules for inhibitory neurons. Middle: boxplot shows changes in composite expression of In-3 module among groups. Right: gene network plot highlights top 25 hub genes. Note that *Mef2c* is among the top 25 hub genes for both excitatory and inhibitory neurons. In boxplots, the center line represents the median, while the lower and upper lines represent the 25th and 75th percentiles, respectively. The whiskers represent the smallest and largest values in the 1.5× interquartile range. Statistical significance values (P) are based on the Kruskal-Wallis test. (F) Representative confocal images of MEF2C protein. Scale bar = 50 μm. (G) Quantification shows MEF2C IHC signals (ANOVA, F (3, 29) = 5.863, P = 0.0029). (H) Western blots of synaptophysin (syn), vGLUT1, MBP, and beta-actin. (I and J) Graphs show expression levels of synaptophysin (I; ANOVA, F (3, 28) = 4.230, P = 0.0138) and vGLUT1 (J; ANOVA, F (3, 28) = 4.371, P = 0.0121). (K) Representative confocal images of vGAT synaptic puncta (scale bar = 20 μm). (L) vGAT positive synaptic puncta was higher in PLX3397+GENUS group (ANOVA, F (3, 29) = 8.831, P = 0.0003). (M, N, O) Confocal images of WFA (scale bar = 100 μm), and 3D rendered WFA and synaptic marker vGLUT1 (scale bar = 20 μm). WFA optical signal (N; ANOVA, F (3, 29) = 4.307, P = 0.0125), vGLUT1 puncta within WFA (O; nested ANOVA, F = 3.300, P = 0.0202). (P) Confocal images of neuronal marker Neun (scale bar = 50 μm). (Q) Density of NeuN in the visual cortex (ANOVA, F (3, 29) = 3.072, p = 0.0433) was significantly increased in the PLX3397 + GENUS group.

To gain additional insights, we adopted a complementary approach involving weighted gene co-expression analysis for E- and I-neurons. The analysis revealed seven and five gene modules for E- and I-neurons, respectively (**Figure 4D and 4E**). Module Ex-2 exhibited the highest effect size and significance when comparing the PLX3397+GENUS to the PLX3397 alone group in E-neurons. Its expression was downregulated in the PLX3397 alone group compared to the control group. However, the co-expression of the module was significantly elevated in the PLX3397+GENUS group (**Figure 4D**). Genes within the Ex-2 module were involved in synapse organization and axonogenesis (**Figure 4D**). Similarly, in I-neurons, the gene module with the most significant effect size and significance comparing the PLX3397+GENUS treatment to the PLX3397 alone group was module In-3, which was enriched for synapse organization and extracellular matrix organization (**Figure 4E**). Compared to the control group, expression of the In-3 module was significantly reduced in the PLX3397 alone group. However, the expression of the module genes was partially restored in the PLX3397+GENUS group. Interestingly, *Mef2c*, a transcription factor, ranked among the top 25 hub genes in both Ex-2 and In-3 gene modules (**Figure 4D and 4E**). Furthermore, *Mef2c* was one of the highest-upregulated genes after combined administration of PLX3397 and GENUS in both E- and I-neurons (**Figure S5**). We observed the upregulation of several MEF2C target genes (**Table S3**) in excitatory (Neat1, Pten, Calm1, Tmsb4x, Slc25a4, Ubc, Ankrd33b, Malat1) and inhibitory (Epb41l2) neurons. Recent findings have demonstrated that MEF2C regulates synaptic genes, intrinsic neuronal functions, and confers resilience to neurodegeneration^19^. To validate this observation based on transcriptomic data, we performed IHC staining of MEF2C to determine its protein expression level. We indeed observed that treatment with PLX3397+GENUS (13.62 ± 0.93) significantly increased the expression of MEF2C compared to control (8.58 ± 0.70), PLX3397 (10.68 ± 0.76), and GENUS (11.08 ± 1.00) alone groups (**Figure 4F and 4G**). Together, these findings are consistent with the notion that repeated GENUS administration in mice treated with PLX3397 improved the expression of genes that impact neural function.

To further validate these findings, we performed additional biochemical analyses. Western blot analysis of synaptic proteins revealed that PLX3397+GENUS administration increased the overall levels of synaptic proteins such as synaptophysin (100 ± 12.03, 185.3 ± 23.83, 142.1 ± 9.22, &190.5 ± 27.31 in control, PLX3397, GENUS, PLX3397+GENUS groups, respectively) and vGLUT1 (100 ± 7.30, 130.1 ± 10.05, 108.7 ± 6.584 & 137.8 ± 9.176) (**Figure 4H-4J**). Interestingly, IHC analysis showed PLX3397 (109.9 ± 2.235), GENUS (108.6 ± 3.01), and PLX3397+GENUS (122.6 ± 4.14) administrations increased vGAT synaptic puncta compared to control (100 ± 2.71) (**Figure 4K and 4L**). While vGAT puncta did not differ between PLX3397 and GENUS groups, we observed a stronger increase in vGAT puncta in 5xFAD mice that received PLX3397+GENUS in the visual cortex (**Figure 4L**), suggesting higher levels of inhibitory synaptic connectivity in 5xFAD mice receiving combined PLX3397+GENUS. Next, we observed that PLX3397 (130.4 ± 7.178), GENUS (116.2 ± 5.225) and PLX3397+GENUS (119.0 ± 6.325) all resulted in increased WFA signals, with the highest levels in the PLX3397 alone group compared to controls (100 ± 4.655) (**Figure 4M and 4N**). Further, WFA content was higher within the microglia in the PLX3397 group but not in PLX3397+GENUS compared to control mice (**Figure S6A-S6C**), suggesting an active role of microglia in organizing PNN in L4, and further that GENUS transforms this microglial phenotype. Synaptic inputs maintained through the structure of PNNs wrapped around PV interneurons are shown to modulate the activity of PV interneurons robustly^45^. We thus performed triple labeling (PV, WFA and vGLUT1) to examine the excitatory presynaptic marker vGLUT1, one of the genes highly impacted by PLX3397+GENUS treatment (by RNA-seq), colocalizing within WFA+ PNNs surrounding PV interneurons. We observed that PLX3397+GENUS (366.6 ± 12.78), & GENUS (363.0 ± 16.64), but not PLX3397 alone (332.7 ± 13.14 versus 313.4 ± 13.20 in controls), significantly increased vGLUT1 synaptic puncta (**Figure 4M and 4O**).

Given these observations, we next assessed neuronal density and found that 5xFAD mice after PLX3397+GENUS but not PLX3397 or GENUS had higher NeuN+ neuronal density in the visual cortex (**Figure 4P and 4Q**), suggesting that combining CSF1R inhibition with GENUS may provide better neuroprotective effects in 5xFAD mice.

Finally, we performed a behavioral analysis to assess whether the increased genes related to learning and memory observed in both excitatory and inhibitory neuronal clusters after PLX3397+GENUS, together with improved neural oscillatory activity and higher neuronal densities, are associated with better cognition. Following treatment with PLX3397, GENUS, and PLX3397+GENUS, 5xFAD mice were tested in an open field (OF), followed by a novel object recognition (NOR) test to assess memory. In the OF test, we did not observe any changes in the time spent in the center of the OF arena nor changes in locomotor activity (**Figure S7A-S7E**). In the NOR test, 5xFAD mice treated with GENUS (60.30 ± 3.58) showed improved recognition memory compared to the chance level (50%). Similarly, PLX3397+GENUS (72.69 ± 3.11) showed an improved preference for the novel object compared to the chance level (50%). Notably, PLX3397+GENUS also showed significantly higher recognition memory than the control (51.33 ± 4.15) group (**Figure S7E**), suggesting that PLX3397+GENUS offers cognitive benefits in AD mice.

## DISCUSSION

Our understanding of the role of microglia in neural circuit function and oscillations is evolving. Oscillations emerge when groups of cells synchronize their transmembrane currents and neuronal spiking. The spiking of many single neurons is synchronized such that they spike at a preferred phase of the oscillations. In particular, theta and gamma oscillations in the visual cortex are involved in attention, learning, and memory^46–49^. In the present study, we describe a previously uncharacterized function of microglia on neural oscillations: 1) in the absence of CSF1R-sensitive microglia in the adult animals, neuronal spiking and theta/gamma oscillations are decoupled, 2) this decoupling is associated with changes in genes related to synapse organization, and 3) repeatedly driving gamma entrainment in PLX3397 treated 5xFAD mice impacted the neuronal transcriptional profile related to improved synaptic integrity, restored neural oscillations, and rescued novel object recognition memory.

L4 neurons in the primary visual cortex (V1) receive robust input from the lateral geniculate nucleus (LGN)^50^. In 5xFAD mice receiving PLX3397, we observed cortical layer-specific neuronal spiking patterns with L4 interneurons showing dramatic reductions while E-neurons increased spiking rate during the onset of aberrant theta activity, indicating abnormal synaptic connectivity and communication between LGN-V1 cortex in 5xFAD mice with reduced microglia. Microglia play an indispensable role in synaptic and circuit organization in adult animals, a role consistent with their involvement in orchestrating LGN-V1 connectivity during development^51,52^. While additional synaptic markers are evident after CSF1R-sensitive microglial removal, L4 PV interneurons show aberrant alterations in their synaptic input architecture, as indicated by changes in PNN and vGLUT1 density. Our data demonstrate that GENUS significantly modifies the synaptic connectivity within PNN surrounding L4 PV interneurons, which is closely associated with improved neural oscillations in PLX3397-treated 5xFAD mice. Further, enhanced synaptic density after microglial removal is thought to be attributed to reduced synaptic pruning by microglia, and this aberrantly regulates neural communications. Additionally, the inhibitory synaptic marker vGAT was also significantly enhanced after PlX3397+GENUS compared to control and PLX3397 alone, suggesting an overall increase in inhibitory synapses. Interestingly, our unbiased gene expression analysis reveals that driving gamma induces intrinsic neuronal mechanisms to enhance the expression of genes related to synaptic functions, extracellular architecture, and myelination in 5xFAD mice. Notably, MEF2C has recently been identified as a resilience factor against neurodegeneration^19^, and is also a regulator of activity-dependent synapse elimination^53^, which is relevant to our findings. Additionally, in conjunction with glia-dependent enhancements in synaptic connectivity, gamma entrainment provides neural circuit protection that counteracts the increase in synaptic density resulting from CSF1R inhibitor-induced microglial depletion.

In PLX3397-treated 5xFAD mice, we observed aberrant theta and gamma oscillatory activities. A more potent CSF1R inhibitor, PLX5622, also impacted neural activities and reduced seizure susceptibility in mice^15^. Importantly, our findings demonstrate that GENUS in the PLX3397-treated mice significantly reduced aberrant theta bursts. Moreover, in human subjects with epileptic discharges in the occipital areas, visual sensory flicker evoked gamma significantly reduced interictal discharges^54^. Thus, although gamma oscillations have long been shown to be involved in higher-order cognitive processes, including attention and memory, these recent findings suggest their value in regulating aberrant oscillations, including in epilepsy. These findings also highlight the need for approaches such as gamma stimulation to regulate neural activities and maintain a healthy cognitive status in people taking PLX3397 to treat TGC.

Consistent with our previous findings^30^, we observed that GENUS reduced microglial density in 5xFAD mice, but it should be noted that the reduction is not as dramatic as PLX3397 administration. Previously, GENUS was thought to act to transform microglia to provide beneficial effects; however, recent observations indicate that chronic GENUS reduces microglial density and inflammatory response^43,44^. Thus, these findings would be consistent with the view that lowering microglia would offer benefits in AD. As microglial functions, specifically their homeostatic ones, are critical, how much reduction of microglia is optimal for the protective effect in AD remains to be thoroughly studied.

We and others have previously shown that GENUS improves synaptic density in multiple models^27,44^ and improves oscillatory activity. While several studies showed that PLX3397 administration also improves synaptic density, the oscillatory activities are somewhat dysregulated. Our results provide evidence that combining PLX3397 and GENUS not only preserves synaptic density but also positively regulates neural oscillatory activity in AD mice.

These findings suggest that anti-inflammatory drugs, such as PLX3397, which show great promise for pathological modification, can be combined with non-invasive sensory stimulation to offer neuroprotection and cognition in AD. Due to the current limits of PLX3397 in stem cell-based therapies, researchers recently engineered an inhibitor-resistant CSF1R variant as a way of ensuring efficient cell replacement of diseased microglia^55^; however, the long-term function of these transplanted microglia is unknown, especially in terms of neuronal oscillatory activity, and as such, combined therapy with gamma entrainment may be beneficial in sustaining a healthy brain environment for engrafted cells. It is thought that despite the persistence of pathological proteins, peripheral macrophages will not enter the brain after microglial depletion^33,56^, highlighting the importance of modifying the brain environment through additional pathways (i.e. neuronal spiking/oscillations) that can prevent newly replaced microglia from adopting a diseased phenotype similar to that of their predecessors. Therefore, our findings provide proof of the principle that a combination of microglial-targeted pharmacology and brain stimulation is a promising strategy to improve AD, and also perhaps tumor treatment outcomes. Although confirming efficacy in human disease will require clinical trials, assessing combinatory PLX3397+GENUS therapy in patients has translational promise given that recent studies have shown the safety and feasibility of GENUS in humans^57–63^, and that the FDA has already approved PLX3397 for the treatment of TGC^17,64^. Remarkably, recent clinical trials of gamma sensory stimulation showed that daily stimulation reduces brain atrophy and increases functional connectivity^59^, which may reflect increased neuronal survival and synaptic integrity, similar to our findings. Recently, microglia have received much interest for their role in driving AD pathogenesis, but microglia-targeted treatments have yet to advance to clinical applications. We showed here that combining microglial reduction with gamma sensory stimulation provided the most robust neuroprotective effects. Thus, our study identifies the role of microglia in modulating brain oscillatory activity and provides evidence supporting the potential of combining microglia-targeted therapies with neural entrainment approaches, such as GENUS, for the treatment of neurodegenerative diseases.

## ACKNOWLEDGMENTS

We thank all the members of the Tsai Laboratory for their valuable comments during many discussions of this work. We specifically thank Rosalind Mott Firenze, Cheng-Yi Yang and Jung Park for carefully reading the manuscript and providing helpful comments. We thank Erica McNamara for animal colony maintenance and Zhuyu Peng for snRNA-seq sample preparation, supporting this work. We are grateful to the funders of this work including the National Institutes of Health grants R56AG069232, RO1 AG069232, and RO1 NS122742 to L-HT; Freedom Together Foundation, The Dolby Family, Robert A. and Renee E. Belfer Family Foundation, Carol and Gene Ludwig Family Foundation, Che King Leo, Lawrence and Deborah Hilibrand, Kathy and Miguel Octavio, Marc Haas Foundation, Dave and Mary Wargo, James D. Cook, Norbert H. Hardner Foundation, and the CBR startup fund — all without which this work would not have happened. The funders had no role in study design, data collection and analysis, the decision to publish or the preparation of the manuscript.

## AUTHOR CONTRIBUTIONS

Conceptualization: A.C, L.-H.T. Methodology: A.C., C.P., Z.P., M.R.I., P.L.B. Investigation: A.C., M.S., P.L.B., N.S, M.R.I, K.A., Visualization: A.C., M.R.I., N.S., Funding acquisition: L.-H.T. Project administration: A.C. L.H.T. Supervision: L.-H.T. Writing—original draft and revision: A.C., M.R.I, P.L.B., and L.-H.T.

## INCLUSION AND DIVERSITY

We worked to ensure sex balance in the selection of non-human subjects. While citing references scientifically relevant for this work, we also actively worked to promote gender balance in our reference list.

## STAR METHODS

### Key Resources Table

| REAGENT or RESOURCE | SOURCE | IDENTIFIER |
| --- | --- | --- |
| **Antibodies** |  |  |
| Anti-IBA1 (1:500) | Synaptic Systems | Cat#234 004 |
| Anti-IBA1 (1:500) | Wako Chemicals | Cat#019-19741 |
| Anti-Parvalbumin (1:500) | Swant | Cat#GP72 |
| Anti-Neun (1:1000) | Synaptic Systems | Cat#266 004 |
| Anti-GFAP (1:500) | Thermo Fisher Scientific | Cat#130300 |
| Anti-MEF2C (1:500) | Cell Signaling Technology | Cat#5030T |
| Anti-Amyloid (D54D2) | Cell Signaling Technology | Cat#8243 |
| Anti-vGAT (1:500) | Synaptic Systems | Cat#131 013 |
| Anti-vGLUT1 (1:500) | Synaptic Systems | Cat#1135 302 |
| Anti-C1q (1:500) | Abcam | Cat#ab182451 |
| Anti-synaptophysin (1:500) | Sigma | Cat#S5768 |
| Anti-MBP (1:500) | Abcam | Cat#ab40390 |
| Biotinylated Wisteria Floribunda lectin | Vector Laboratories | Cat#B-1355 |
| Donkey Anti-Rabbit, Alexa Fluor® 488 conjugate | Invitrogen | Cat#A-21206 |
| Donkey Anti-Mouse, Alexa Fluor 488 conjugate | Invitrogen | Cat# A21202 |
| Donkey Anti-Rabbit, Alexa Fluor 594 | Invitrogen | Cat# A11037 |
| Donkey Anti-Goat, Alexa Fluor 594 | Invitrogen | Cat# A11058 |
| Donkey Anti-Mouse, Alexa Fluor 594 | Invitrogen | Cat# A11032 |
| Donkey Anti-Guinea Pig, Alexa Fluor 647 | Invitrogen | Cat#A-21450 |
| Donkey Anti-Rabbit, Alexa Fluor 647 | Invitrogen | Cat#A-31573 |
| Donkey Anti-Mouse, Alexa Fluor 647 | Invitrogen | Cat#A-31571 |
| Streptavidin Conjugated Alexa Fluor 594 | Thermo Fisher Scientific | Cat#S32356 |
| Horseradish peroxidase-linked secondary antibody | Jackson Immuno Research | Cat#211-032-171 |
| **Chemicals and Kits** |  |  |
| PLX3397 (pexidartinib) | Medko | CAS#1029044-16-3 |
| Western-Bright Quantum Kit | Advansta | Cat#K-12042-D20 |
| Laemli Sample Buffer | Bio-Rad | Cat#1610747 |
| Premade polyacrylamide Gels | Bio-Rad | Cat#4561096 and 4561094 |
| Chromium Next GEM Single Cell 3' Kit v3.1 | 10X Genomics | Cat#PN-1000268 |
| Dental Cement, C & B Metabond quick adhesive | Parkell | Cat#836SKU:S380 |
| **Experimental animals** |  |  |
| 5xFAD (B6.Cg-Tg(APPswFLon, PSEN1*<sup>M146L</sup>*<sup>L286V</sup>)6799Vas/Mmj ax) | Jackson Laboratory | MMRRC Stock#34848-JAX |
| B6;CBA-Tg(Camk2a-tTA)1Mmay/J | Jackson Laboratory | Stock No: 003010 |
| C57BL/6-Tg(tetO-CDK5R1/GFP)337Lht/J | Jackson Laboratory | Stock No: 005706 |
| **Software and Algorithms** |  |  |
| Imarisx64 | Bitplane | Version 9.3 & Version 9.7.1 |
| ImageJ | NIH | Version 1.52a |
| Zen acquisition program | Zeiss | N/A |
| Neuralynx Matlab Import Export | Neuralynx | Version 6.0.0 |
| GraphPad Prism | GraphPad | Version 8.3.1 & Version 9.3.1 |
| Matlab | MathWorks | Version 2019a |
| MATLAB File Exchange: Circular Statistics Toolbox | MATLAB File Exchange Function<br>(https://www.mathworks.com/matlabcentral/fileexchange/10676-circular-statistics-toolbox-directional-statistics) | N/A |
| EthoVision XT | Noldus | Version XT 15 & Version XT 17 |
| Chem doc | Bio-Rad | N/A |
| CellRanger | 10x Genomics | V3.0 |
| Seurat | R (CRAN/Bioconductor) | v 4.1.0 |
| WGCNA | R package (CRAN) | v 1.70-3 |
| clusterProfiler | R package (Bioconductor) | v 4.2.0 |
| rrvgo | R package (Bioconductor) | v 1.6.0 |

## CONTACT FOR REAGENT AND RESOURCE SHARING

All data necessary to assess the conclusions of this research are available in the text and supplementary materials. Further information and requests for resources and reagents should be directed to the Lead Contact, Li-Huei Tsai.

## ANIMAL MODELS

All the experiments were approved by the Committee for Animal Care of the Division of Comparative Medicine at the Massachusetts Institute of Technology (MIT) and carried out at MIT. Tg(Camk2a-tTA), and Tg(APPSwFlLon,PSEN1*M146L*L286V) were obtained from the Jackson laboratory. Tg(tetO-CDK5R1/GFP) was generated in our lab. 5xFAD mice were 10-12 months old and CK-p25 mice were 8months-old prior to commencement of experiments. Equal numbers of male and female CK-p25 were used in each group. Female 5xFAD mice were used for the RNA-sequencing experiment, and male 5xFAD mice were used for all other experiments. We used equal numbers of male (n = 4 - 6 mice per group) and female (n = 4 – 6 mice/group) CK-p25 mice for all experiments.

### Experimental treatment

CSF1R inhibitor PLX3397: We obtained PLX3397 (Pexidartinib; CAS#: 1029044-16-3, medkoo.com/products/4501) drug from Medkoo Biosciences (Morrisville, NC, USA). PLX3397 was then irradiated and premixed into the rodent diet at 600 ppm (PMI RMH 3000 5P76 rodent diet with 0.06% PLX3397). A red food color was added to the PLX3397 diet. These subsequent processes were completed by Envigo Teklad Diets (Madison, WI, USA). PLX3397 diet was stored in a cold room until use.

PLX3397 administration: Mice were introduced into clean new cages, and the regular diet was replaced with a diet containing PLX3397. Only experimenters A.C, M.S, and C.P handled or changed cages during the entire experimental procedure. Cages were changed once weekly. Mice were given PLX3397 diet and water ad libitum, just as the regular diet control mice throughout the experiment.

GENUS stimulation: Light flicker stimulation was delivered as previously described^30,65^. Mice were transported from the holding room to the flicker room on adjacent floors of the same building. Mice were habituated under dim light for 20 min before the start of the experiment, and then introduced to the stimulation cage (similar to the home cage, except without bedding and three of its sides covered with black sheeting). All GENUS protocols were administered on a daily basis for 1h/d for the number of days as specified. Mice were allowed to freely move inside the cage but did not have access to food or water during the 1 h light flicker stimulation. An array of light-emitting diodes (LEDs) was present on the open side of the cage and was driven to flicker at a frequency of 40 Hz with a square wave current pattern using an Arduino system. The luminescence intensity of light that covered the total area of GENUS stimulation cage varied from ∼200 - 1000 lux as measured from the back and front of the cage (mice were free to move within the cage). After 1h of light flicker exposure, mice were returned to their home cage and allowed to rest for a further 30 min before being transported back to the holding room. No-stimulation control mice underwent the same transport and were exposed to similar cages with similar food and water restriction in the same room, but experienced only normal room light for the 1 h. The experimenters who stimulated the mice were male.

#### Experimental group description

Control 5xFAD mice: Mice received a regular rodent diet and water ad libitum. Mice also received control sensory stimulation as described above.

PLX3397 5xFAD mice: Mice received PLX3397 and water ad libitum for 50 days. GENUS 5xFAD mice: Mice were subjected to 30 days of daily GENUS (1h/d).

PLX3397+GENUS 5xFAD mice: Following 20 days of administration of PLX3397 chow, 5xFAD mice were subjected to 30 days of daily GENUS (1h/d). Mice were still maintained on the PLX3397 diet during the 30 days of GENUS stimulation.

Control CK-p25 mice: p25 was induced by replacing the doxycycline diet with a regular rodent diet. Mice also received control sensory stimulation as described above. This treatment procedure (regular diet + control stimulation) was administered for 6 weeks.

PLX3397 CK-p25 mice: p25 was induced by replacing the doxycycline diet with a PLX3397-containing diet. This treatment procedure (PLX3397 diet) was administered for 6 weeks.

GENUS CK-p25 mice: p25 was induced by replacing the doxycycline diet with a regular rodent diet. Mice also received 40 Hz sensory stimulation as described above. This treatment procedure (regular diet + 40 Hz stimulation) was administered for 6 weeks.

PLX3397+GENUS CK-p25 mice: p25 was induced while also inhibiting CSF1R by replacing the doxycycline diet with the PLX3397 rodent diet. In addition, CK-p25 mice were also subjected to daily GENUS (1h/d) for 6 weeks simultaneously.

### Open Field (OF) and novel object recognition (NOR) test

For OF, mice were introduced into an open field box (dimensions: length = 460mm, width = 460mm and height = 400mm; TSE-Systems) and were tracked using Noldus (Ethovision) for 12 min, with time spent in the center and peripheral area of the arena measured. NOR test was conducted on the following day, when mice were re-introduced into the same open field box which now additionally contained two identical novel objects, and were allowed to explore the objects for 7 min (novel object habituation). Mice were then placed back in their home cages for 20 min after the last exploration. They were then returned to the same arena, with one of the two objects replaced with a new object. Mouse behavior was monitored for 7 min. Time spent exploring both the familiar and novel objects was recorded using Noldus and computed offline. The percentage of novelty preference index was calculated as follows: time exploring novel object (Nt) divided by total time exploring novel and familiar (Ft) objects and presented in %-{[Nt/Nt+Ft]*100}. All 5xFAD mice used were age-matched males. Age-matched 4-6 male and 4-6 female mice/group were used from the CK-p25 mice cohort.

## IMMUNOHISTOCHEMISTRY

Mice were transcardially perfused with 40 mL of ice-cold phosphate-buffered saline (PBS) followed by 40 mL of 4% paraformaldehyde (PFA; Electron Microscopy Sciences, Cat#15714-S) in PBS. Brains were removed and post-fixed in 4% PFA overnight at 4°C and transferred to PBS before sectioning. Brains were mounted on a vibratome stage (Leica VT1000S) using superglue and sliced into 40 μm sections. Slices were subsequently washed with PBS and blocked using 5% normal donkey serum prepared in PBS containing 0.3% Triton X-100 (PBST) for 2 h at room temperature. The blocking buffer was aspirated out and the slices were incubated with the appropriate primary antibody (prepared in fresh blocking buffer) overnight at 4°C on a shaker. Slices were then washed three times (10 min each) with the blocking buffer and then incubated with Alexa Fluor 488, 555, 594 or 647 conjugated secondary antibodies for 2 h at room temperature. Following three washes (15 min each) with blocking buffer and one final wash with PBS (10 min), slices were mounted with fluromount-G (Electron microscopic Sciences).

Antibodies: IBA1 (Synaptic Systems, Cat # 234 004, dilution-1:500; Wako Chemicals, Cat # 019-19741, dilution-1:500), GFAP (Thermo Fisher Scientific, Cat # 130300, dilution-1:500), MEF2C (Cell Signaling Technology, Cat # 5030T), D54D2 (CST, Cat # 8243, dilution-1:500), vGAT (Synaptic Systems, Cat # 131 013, dilution-1:500), vGLUT (Synaptic Systems, Cat # 1135 302, dilution-1:500), NeuN (Synaptic Systems, Cat # 266 004, dilution-1:1000), C1q (Abcam, Cat # ab182451, dilution-1:500), synaptophysin (Sigma, Cat # S5768). The following combination of secondary antibodies were used: (1) Alexa Fluor 488, 594 and 647, (2) Alexa Fluor 555 and 647, (3) Alexa Fluor 594 and 647, or (4) Alexa Fluor 488 and 647. All secondary antibodies were obtained from Invitrogen. Biotinylated Wisteria Floribunda Lectin (Vector Laboratories, Cat # B-1355, dilution-1:500) followed by streptavidin conjugated Alexa Fluor 594 (Thermo Fisher Scientific, Cat# S32356, dilution-1:1000) was used to examine WFA.

Images were acquired using either LSM 710 or LSM 880 confocal microscopes (Zeiss) with 10x, 20x, or 40x objectives at identical settings for all conditions. Images were quantified using Imarisx64 9.3 or Imarisx64 9.7 (Bitplane, Switzerland). For each experimental condition, two coronal sections per mouse from the indicated number of animals were used. The averaged values from the two to four images per mouse were used for quantification. The experimenter blinded to the treatment conditions performed all the image processing and quantification.

C1q and MBP signal intensity: Using an LSM 710 with a 20x or 40x objectives, z stacks of the entire slice thickness 40 µm (40 images from each field) were acquired. The signal intensity was measured in Imaris.

Microglia: Iba1 immunoreactive cells were considered microglia. Using an LSM 710 or LSM 880 with a 10x (for Iba1+ cell counts) or 40x (for morphological analysis) objective z stacks of the entire slice thickness 40 μm with 0.5 μm step size were acquired. Imaris was used for 3D rendering of images to quantify the total volume of microglia.

MEF2C: LSM 710, with a 40x objective, was used to acquire the images. An entire 40 μm thickness of the slices was acquired in Z stacks 40 per image. MEF2C optical signal was measured using Image J.

NeuN positive cell: All images were acquired in Z stacks-10 per image and were quantified. The spot-count inbuilt function in multi-point tool in Imarisx64 9.3 was used to count cells automatically.

vGAT and vGLUT1 puncta: LSM 710, with a 40x objective, was used to acquire the images. An entire 40 μm thickness of the slices was acquired in Z stacks-80 per image (step of 0.5μm). The spot-count inbuilt function in Imarisx64 9.3 (cohort1) and 9.7 (cohort 2) was used to count cells automatically.

Myelinated PV axons: Triple-labeled brain sections with WFA, PV and MBP were used for this analysis. WFA positive PV axons (double positive) were first traced and MBP signal only on those WFA: PV++ axons were visualized using Imaris and the maximum length of that triple positive signal was quantified.

### Western blotting

Brains were perfused with PBS and fixed with 4% PFA. The visual cortex was dissected out into 1.5ml Eppendorf tube containing 100 μl of TS buffer (600mM Tris-HCl, pH 8, and 2% SDS). The tissue was homogenized thoroughly using a handheld gun. The homogenate was incubated at 90 degrees C for 2 h (at 500 rpm in TS buffer). The homogenate was then centrifuged at 1000g for 1min at room temperature, and the upper 60 ul of the sample was transferred to a new Eppendorf tube. Laemmli sample buffer (Bio-Rad, Cat # 1610747) was added to the sample. Samples were loaded onto 4-20 % polyacrylamide gels (Bio-Rad, Cat # 4561096 or 4561094) and electrophoresed (Bio-Rad). Protein was transferred from acrylamide gels to nitrocellulose membranes for 12 min (Semi-dry system, Bio-Rad). Membranes were blocked using BSA (5% w/v) diluted in TBS containing 0.1% Tween-20 (Sigma-Aldrich, Cat # P9416) (TBSTw), then incubated in primary antibodies overnight at 4°C. The following day, they were washed three times with TBSTw and incubated with horseradish peroxidase-linked secondary antibodies (Jackson Immuno Research, Cat # 211-032-171, dilution-1:5000) at room temperature for 2 h. After three further washes with TBSTw, membranes were treated with chemiluminescence substrates Western-Bright Quantum kit (Advansta, Cat # K-12042-D20) and the blots were visualized (Chem doc, Bio-Rad). Signal intensities were quantified using ImageJ 1.46q and normalized to values of loading control.

## *IN VIVO* ELECTROPHYSIOLOGY

Mice were anesthetized with isoflurane, restrained in a stereotactic apparatus and craniotomies were made exposing the visual cortex. Specifically, a 2 x 2 mm piece of skull was removed using a dental drill, which was above the V1 (stereotaxic coordinates relative to bregma; AP −3.2; ML ± 2.5); during this entire procedure, the dura was kept intact and moist with saline. Following the skull removal from above both the right V1, we made two additional drilling holes above the frontal cortex and placed two skull screws. Recording probes (Neuronexus, Cat # A1×32-5mm-25-177-CM32, or A1×16-3mm-50-177-CM16LP) were then fitted to the stereotactic apparatus and aligned to the craniotomy and slowly lowered to ∼50 μm above the cortical target depth. The probe was grounded to a skull screw above the cerebellum. Petroleum jelly (Vaseline, 100% white petrolatum) was gently applied on the cranial window without touching the probe/electrodes, which protected both the brain and probe/electrode. Next, we further lowered the probe and adjusted it to reach the target depth. Finally, the probe was cemented on the skull with dental cement, first with a metabond (Parkell, C&B Metabond Quick Adhesive Cement System, # 836 SKU:S380) followed by a dental cement from Steolting (# 51459). Mice were allowed to recover for a period of 4 days.

Following a 2-3-day habituation period for the recording, recordings commenced with the animals allowed to move freely in their home cages. Data were acquired using Neuralynx SX system (Neuralynx, Bozeman, MT, USA) and signals were sampled at 32,000 Hz. The position of animals was tracked using red light-emitting diodes affixed to the probes. After the experiment, mice underwent terminal anesthesia and electrode positions were marked by electrolytic lesioning of brain tissue with 50 mA current for 10 s through each electrode individually, to confirm their anatomical location.

Spikes: Single units were manually isolated by drawing cluster boundaries around the 3D projection of the recorded spikes, presented in SpikeSort3D software (Neuralynx). Cells were considered pyramidal neurons if the mean spike peak-to-trough length exceeded 220 µs and had a higher peak-to-trough ratio. The extracellular spike waveforms shown in Figs. 1 & 2 are inverted for representative purposes only.

Data analyses: LFPs were first filtered to the Nyquist frequency of the target sampling rate then down-sampled to 1000 Hz. Power spectral analyses were performed using the pwelch function in MATLAB using a 500 ms time window with a 50% overlap.

Time-frequency representation of LFP: The LFP data were downsampled to 1,000 Hz. For the calculation of the wavelet power spectrum, the continuous wavelet transforms (CWT) were applied to the LFP using complex Morlet wavelets returning amplitudes at 226 intervals between 1-100 Hz. CWT based wavelet power spectrum was shown in Fig. 1j, & Extended Data Fig. 1a,b, 2a. For visualizing 40 Hz entrainment at finer frequency resolution (Fig. 2f, & Extended Data Fig. 2b), we used multitaper spectral analysis using Chronux toolbox (chronux.org).

Single unit - LFP phase locking: The relationship between spike times and LFP gamma phase was calculated by mean resultant length using the Circular Statistics Toolbox MATLAB File Exchange Function (https://www.mathworks.com/matlabcentral/fileexchange/10676-circular-statistics-toolboxdirectional-statistics)^66^. Briefly, spikes were sorted and LFP traces were filtered using the continuous wavelet transform returning the instantaneous signal phase and amplitudes. Spike times were linearly interpolated to determine phase, with peaks and troughs of gamma defined as 0 and ±pi radians respectively. The resulting phase values were binned to generate spiking probabilities, for each 20-degree interval. Cells were considered to be phase-locked only if they had a distribution significantly different from uniform (P < 0.05 circular Rayleigh test), with the strength of phase-locking calculated as the mean resultant length. All analyses were performed using MATLAB. All *in vivo* electrophysiological analyses were conducted in MATLAB (Mathworks, #R2019a) utilizing signal processing and image processing toolboxes.

Power spectral density (PSD) of single units: We calculated the interspike interval (ISI) and the spectral power estimate to find the frequency at which the neurons are spiking. To estimate the spike PSD, the spike times were converted into a spike train with a frequency bin of 1 ms. The PSD for this spike train was then calculated by using the MATLAB pwelch function without any windowing operations. The considered peaks with 2SD above the mean in the PSD represent the dominant/rhythmic frequency(ies) at which the neurons are spiking. MATLAB functions can be found here https://github.com/ChinnakkaruppanAdaikkan/Rhythmicity-of-Spike-Trains

## SINGLE CELL RNA SEQUENCING

Mouse primary visual cortices from 11-months old 5xFAD mice were used for single nuclei RNAseq experiment. Frozen brain tissues were subjected to a nuclei extraction experimental protocol adapted from previous study^6^. Briefly, cortices were homogenized using a Wheaton Dounce tissue grinder. The homogenized tissue was filtered through a cell strainer (40-μm), mixed with a working solution and loaded on top of an OptiPrep density gradient to separate nuclei by ultracentrifugation using an SW32 rotor. Nuclei were then collected from the 29%/35% interphase of the density gradient, washed once with PBS containing 0.04% BSA, centrifuged at 300g for 3 min (4 °C). Nuclei were then again washed with 1 ml of PBS containing 1% BSA, and counted using a hemocytometer. Single nuclei RNA libraries were prepared using the Chromium Next GEM Single Cell 3ʹ Kit v3.1 according to the manufacturer’s protocol (10x Genomics). The generated snRNA-seq libraries were sequenced using NovaSeq 6000 at the MIT BioMicro Center. Sequencing raw reads were mapped to the mouse genome (mm10) and the cell-wise gene counts were generated using CellRanger software (v3.0, 10x Genomics). All downstream processing and analyses were performed using Seurat (v 4.1.0)^49^. Cells with low quality as quantified based on several cell quality criteria (e.g., number of expressed genes and unique molecular identifiers (UMI), percentage of reads mapped to mitochondrial genes, presence of doublets, nuclei expressing multiple cell type specific markers) were filtered out and nuclei with high quality transcriptional profiles were retained. Specifically, nuclei that have unique feature counts over 4000 or less than 200 and have >10% mitochondrial counts were discarded and DoubletFinder was used to find and remove potential doublets. Data were normalized to library size, scaled, integrated and top 2000 highly variable genes were used to perform an unsupervised Principal Component Analysis (PCA). For non-linear dimension reduction, Uniform Manifold Approximation and Projection (UMAP) was used employing the first 15 components. Cell type specific marker genes were determined using FindAllMarkers, while the differentially expressed genes were determined using FindMarkers function in Seurat. Genes with FDR<0.05 and abs(avg_log2Fc)>=0.1 were considered as differentially expressed. Similar approach was also applied for analyzing the single cell RNAseq data. Weighted gene co-expression analysis for excitatory and inhibitory neuronal snRNAseq data was done using a modified WGCNA approach as described previously^50^. Briefly, metacells were constructed from excitatory and inhibitory neurons separately based on the mean expression from 25 neighboring cells using the *k*-nearest neighbors. Next, a signed WGCNA approach^51^ was implemented by calculating the component-wise values for topological overlap. Modules with a minimum module size of 25 genes, a deepSplit score of 4 were analyzed and modules with a correlation of greater than 0.8 were merged together. Intra-modular connectivity (kME) of WGCNA (v 1.70-3) was used to define the top 25 Hub genes. Gene ontology analysis was done using clusterProfiler (v 4.2.0) and rrvgo package (v 1.6.0) was used to find and reduce the redundancy of gene ontology terms based on semantic similarity. Additional gene ontology analyses were performed using Metascape^67^.

## STATISTICS AND REPRODUCIBILITY

All IHC and behavioral experiments were blinded. No data were excluded for analysis. All IHC experiments were replicated in two independent experiments of at least 3 mice per group in each experiment, and both replications were successful. For all representative images shown, the images are representative of at least two independent stainings and experiments. Statistical tests and significance for each experiment were calculated as noted in the appropriate figure legends, using t-test, Mann-Whitney test, or one-way ANOVA with a Two-stage linear step-up procedure of Benjamini, Krieger and Yekutieli post hoc analysis. Statistical significance was set at 0.05. Statistical analysis was conducted using Prism (version 9.3 and 9.7.1, GraphPad Software).

## DATA AVAILABILITY

snRNA-seq data will be available from the Gene Expression Omnibus upon publication. All data are available in the main text and the Supplementary Information files. Data analyses have been performed using previously published packages that are available from CRAN or Bioconductor.

## SUPPLEMENTARY INFORMATION

**Figure S1.**
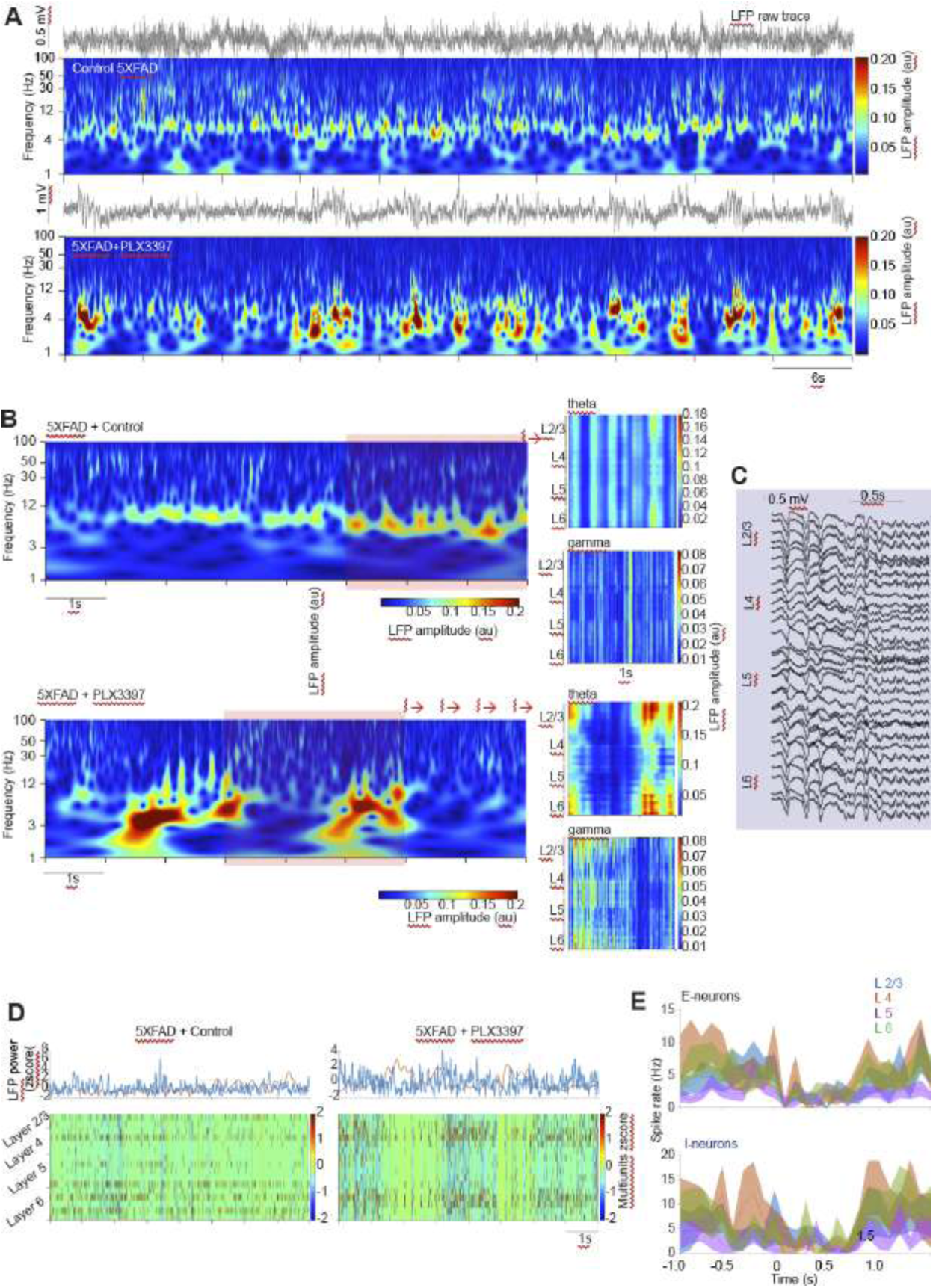
CSF1R-sensitive microglia elimination impacts neural synchrony in 5xFAD mice. (A) Plots show unprocessed raw LFP traces and the corresponding time-resolved power spectra from 5xFAD without or with PLX3397 administration. (B) Power spectrogram from layer 4 LFP (left), and LFP theta or gamma power spectra organized according to cortical depth from control and PLX3397-treated 5xFAD mice. L2/3, L4, L5, & L6 indicates cortical layers 2/3, 4, 5, & 6, respectively. (C) Unprocessed raw LFP traces in PLX3397 5xFAD mice. L2/3, L4, L5, & L6 indicates cortical layers 2/3, 4, 5, & 6, respectively. PLX3397 (D) Top: plots show mean theta (3-12 Hz) and gamma (30-50 Hz) power in 5xFAD without or with Plx3397PLX3397 administration. Multiunit activity is shown at the bottom. (E) Line plots show the mean (± s.e.m) spike rate of E-neurons (top) and I-neurons (bottom) pre-, during-, and post-high theta activity from L2/3, L4, L5, and L6. Time zero represents high theta activity onset.

**Figure S2.**
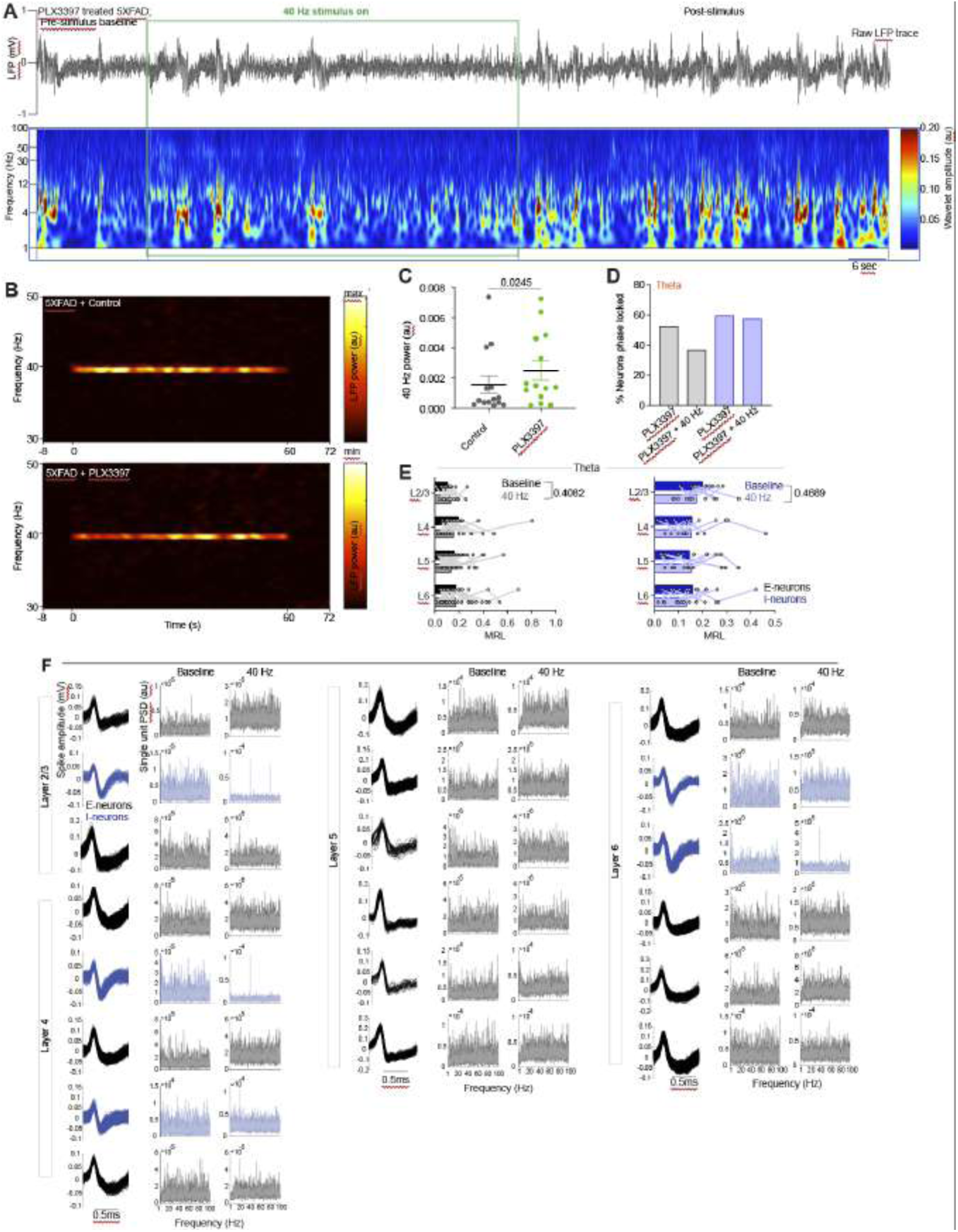
Sensory evoked gamma oscillations in PLX3397 treated 5xFAD mice. (A) LFP trace (top) during pre-stimulation in PLX3397-treated 5xFAD mice — the corresponding wavelet LFP spectrogram before, during, and after acute 40 Hz stimulation. Note the reduction in theta activity during the 40 Hz entrainment. (B) LFP power spectrum in 5xFAD mice with or without PLX3397 administration for 50 days. PLX3397 administered 5xFAD mice exhibited clear 40 Hz entrainment during acute 60sec stimulation. (C) The summary graph shows the mean power of 40 Hz entrainment in control and PLX3397 administered 5xFAD mice. (D) Plot shows the percentage of total neurons phase-locked to theta oscillations based on circular Rayleigh statistics. (E) Plots show the strength of phase locking between neuronal spiking and LFP theta. No significant effect was observed in E-neurons (F (1, 97) = 0.6899, P = 0.4082) and I-neurons (ANOVA, F (1, 43) = 0.4873, P = 0.4889) between baseline and 40 Hz entrainment. (F) Twenty simultaneously recorded single units are organized according to cortical layers in PLX3397+GENUS-treated mice. Spike waveforms of isolated units and the power spectral density of units are shown.

**Figure S3.**
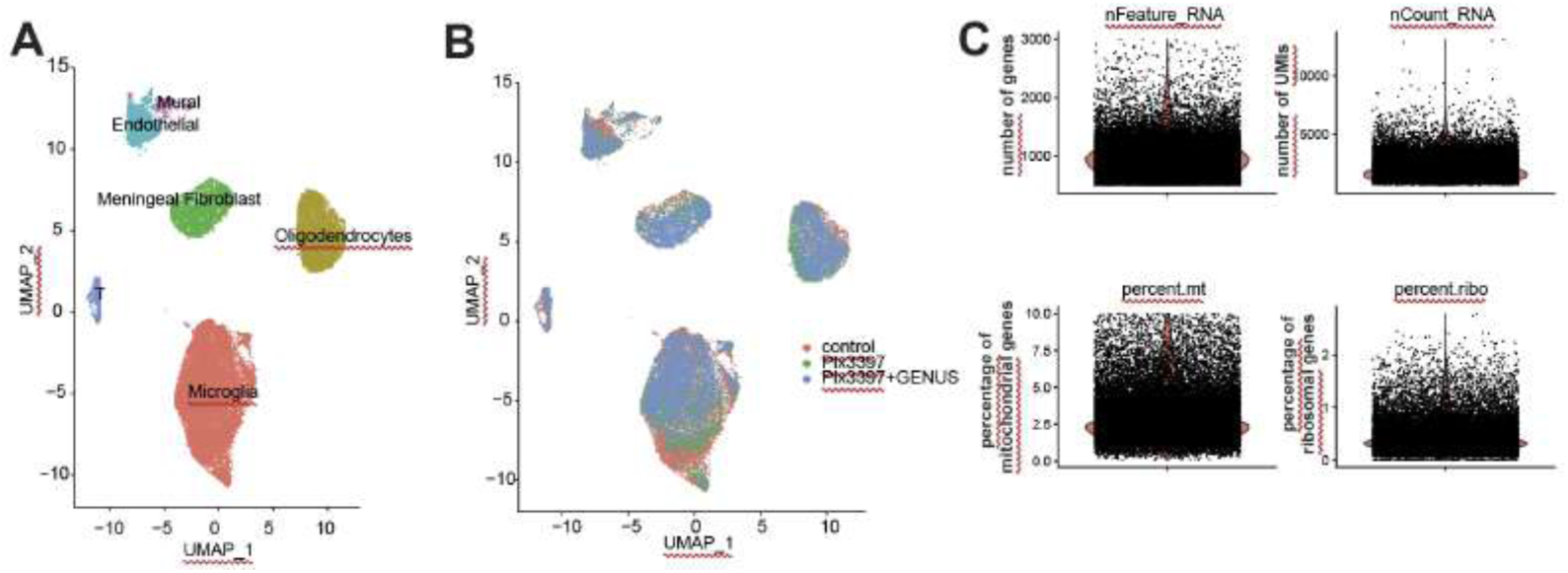
Single-cell transcriptomics profiling from the visual cortices of 5xFAD mice. (A) UMAP plot displaying cell clusters corresponding to microglia, oligodendrocytes, meningeal fibroblasts, endothelial cells, mural cells, and T cells. (B) UMAP plot demonstrating that the cell clusters are well integrated across different treatment conditions. (C) Violin plots showing the distribution of quality control metrics, including the number of detected genes, UMIs, and the percentage of mitochondrial and ribosomal gene expression.

**Figure S4.**
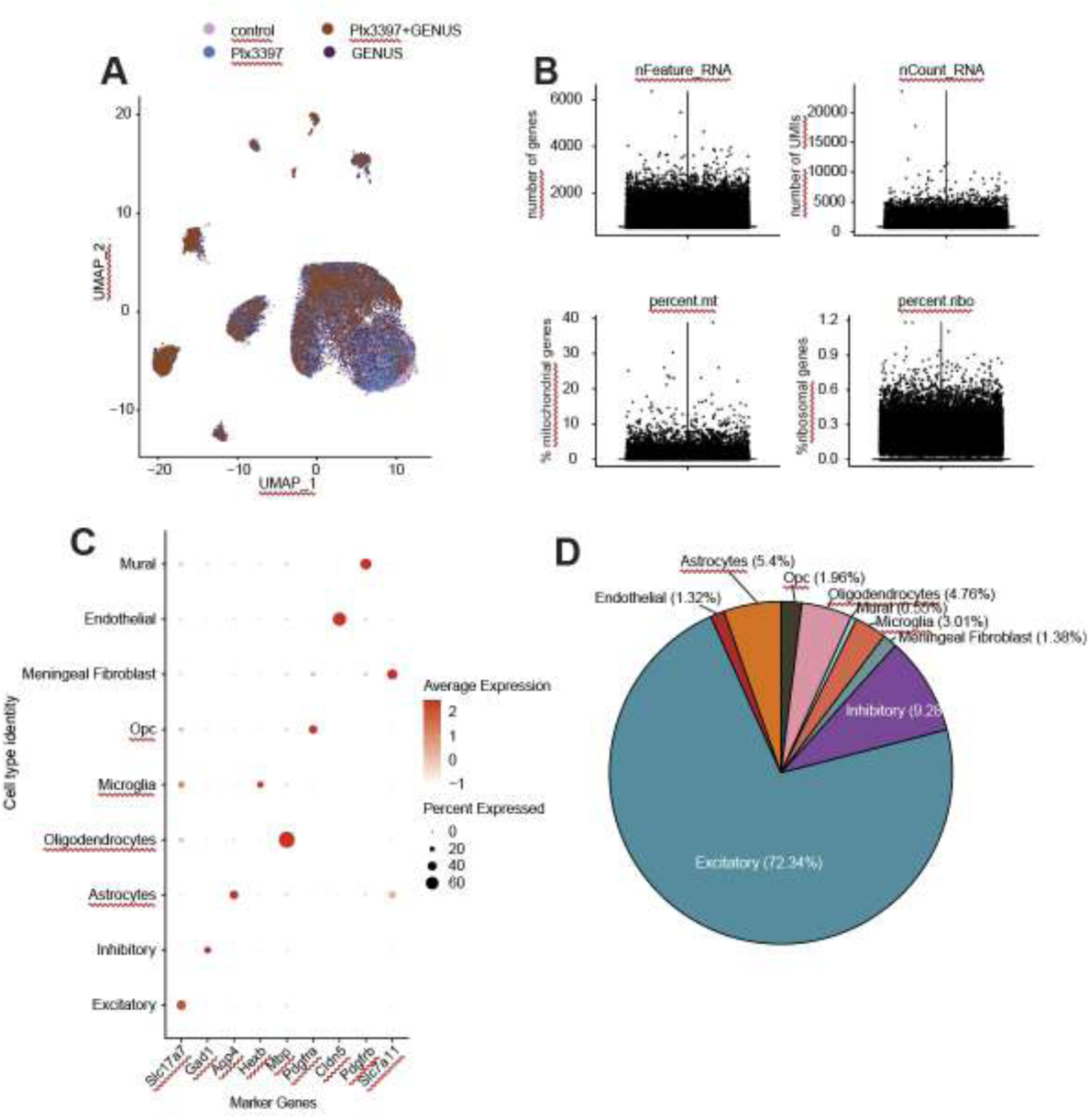
Single-nuclei transcriptomics profiling from the visual cortices of 5xFAD mice. (A) UMAP plot demonstrating well-integrated cell clusters across different treatment conditions. (B) Violin plots depicting the distribution of quality control metrics, including the number of detected genes, UMIs, and the percentage of mitochondrial and ribosomal gene expression. (C) Dot plots illustrating the expression of cell-type-specific marker genes within their respective cell types. (D) Pie chart showing the proportional distribution of identified cell types analyzed in the dataset.

**Figure S5.**
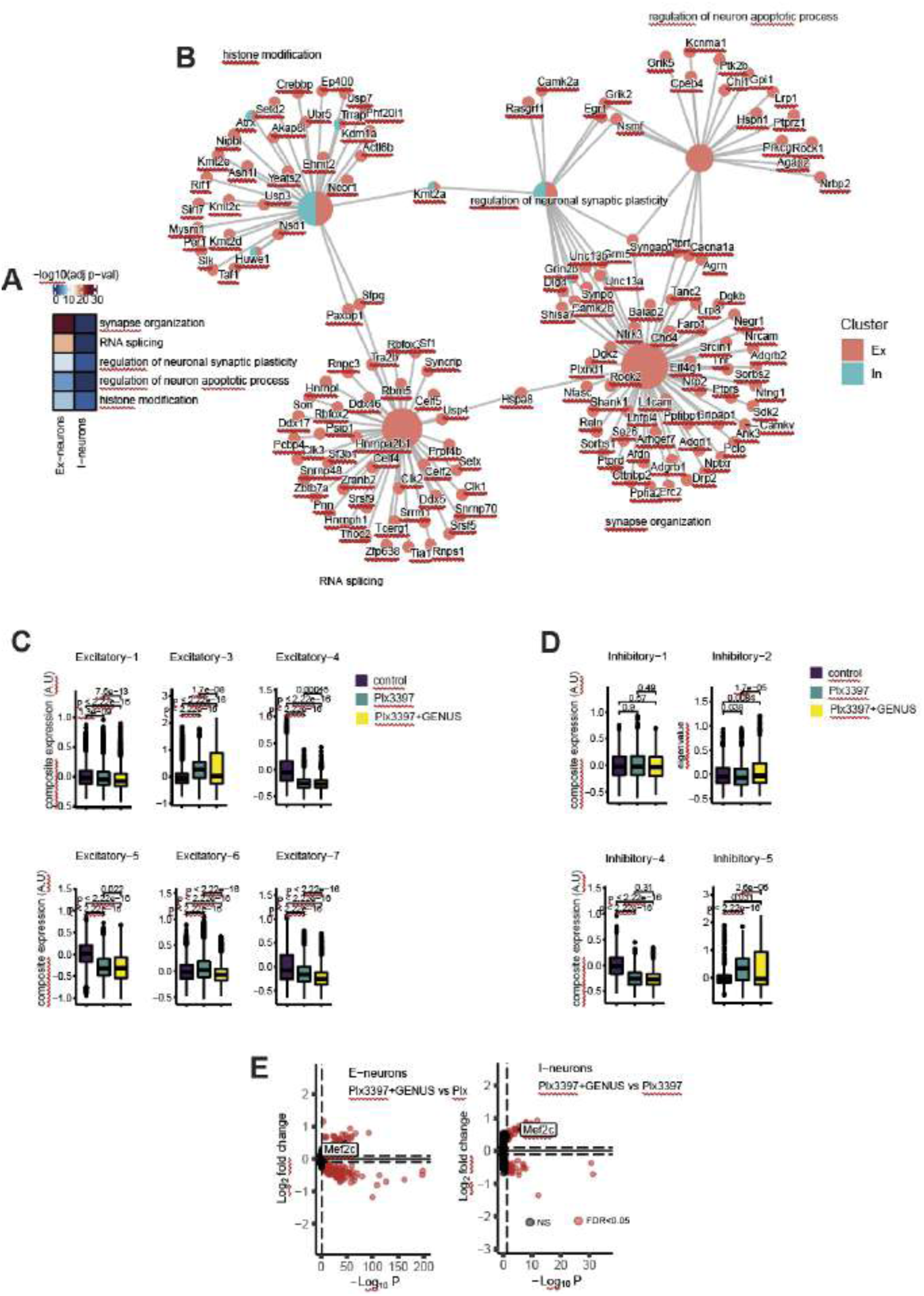
Administration of chronic gamma entrainment enhances Mef2c in both E- and I-neurons of PLX3397 treated 5xFAD mice. (A, B) Gene ontology (GO) analysis for the downregulated genes comparing PLX3397+GENUS and PLX3397 alone in E- and I-neurons. (A) Top GO terms represented in E- and I-neurons for the downregulated genes. (B) Gene network with nodes and edges highlighting the genes involved in the corresponding GO term. Larger nodes represent the gene ontology term while the smaller nodes highlight the genes. Red and green colors represent the observations in the E- and I-neurons respectively. (C, D) Expression of gene modules for (C) excitatory and (D) inhibitory neurons among experimental groups. The midline of the Boxplots indicates the median, while the lower and upper lines represent the 25th and 75th percentiles, respectively. The whiskers represent the smallest and largest values in the 1.5× interquartile range. Kruskal-Wallis test. (E) Volcano plots highlight Mef2c as significantly upregulated in both E- and I-neurons.

**Figure S6.**
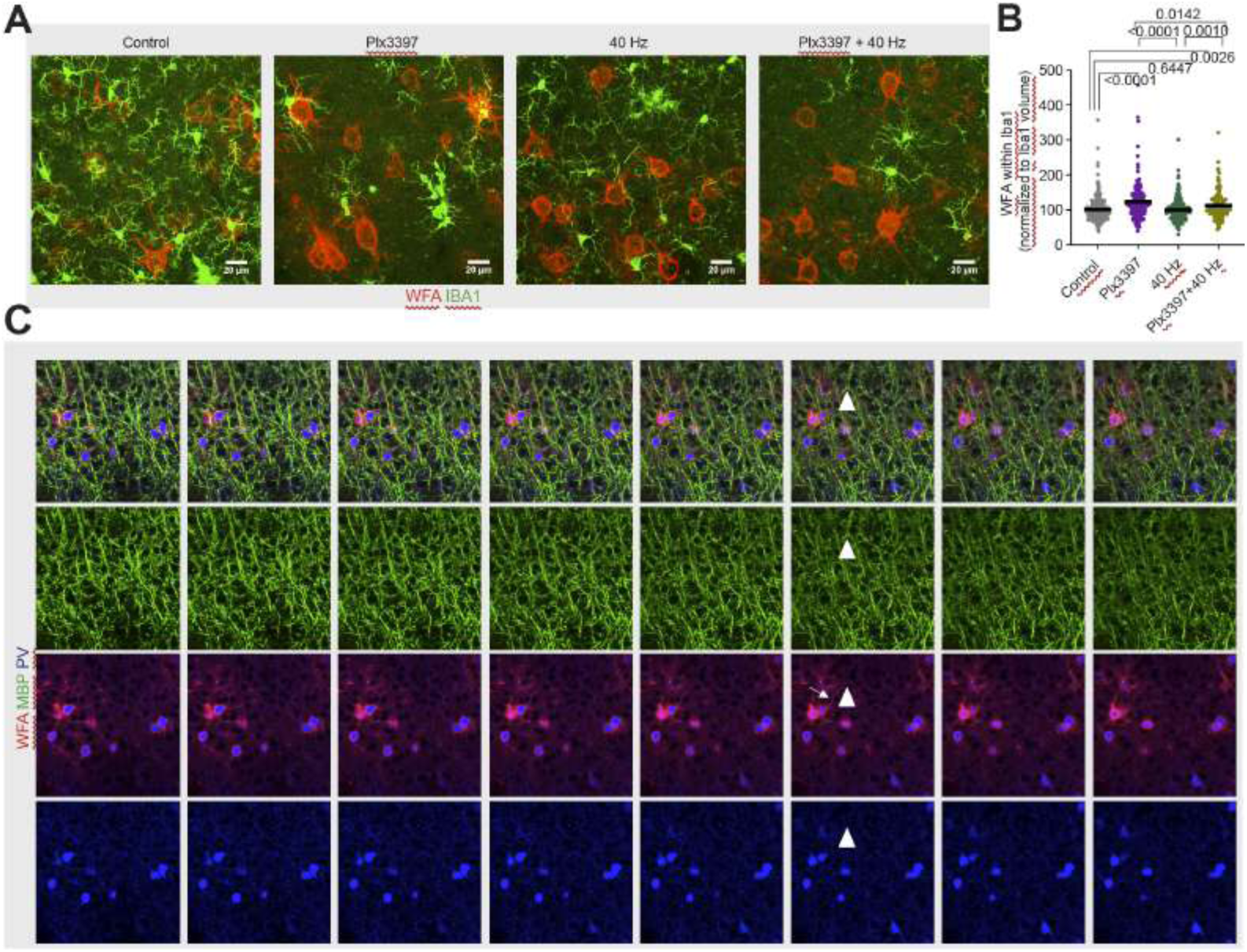
Chronic administration of PLX3397 and GENUS perineuronal net. (A) Representative confocal images show WFA and IBA1 (scale bar = 20 μm). (B) Plot shows the WFA signal within IBA1 (ANOVA F= 17.96, p < 0.0001). (C) Representative serial single-plane confocal images show WFA, MBP, and PV. Note that MBP signals around the axonal process of PV interneurons are evident immediately after WFA but not within WFA. This suggests a multifaceted regulation of the PV axon through myelination and PNN.

**Figure S7.**
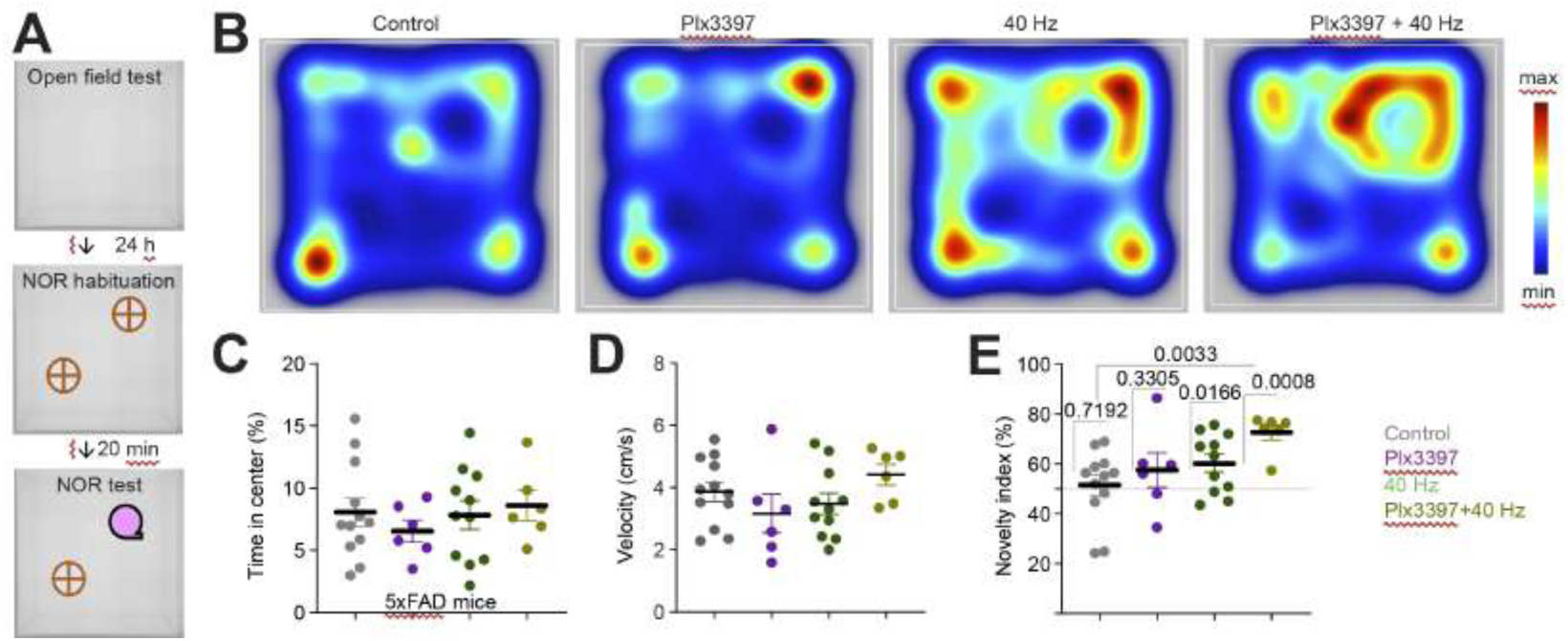
Chronic administration of PLX3397 and GENUS improves novel object recognition memory in 5xFAD mice. (A) Schematic of OF and NOR test. (B) Mice occupancy heatmaps during the NOR test in 5xFAD mice. (C) Time spent in the center during OF (c, ANOVA, F (3, 31) = 0.3847, P = 0.7647) did not differ between groups. (D) Plot shows the velocity of mice during a novel object recognition memory test in 5xFAD mice (ANOVA F (3, 31) = 1.437, P = 0.2508. (E) Novelty index during NOR test was higher in PLX3397+GENUS-treated 5xFAD mice (ANOVA, F (3, 31) = 3.456, P = 0.0282).

